# DEK influences the trade-off between growth and arrest via H2A.Z-nucleosomes in Arabidopsis

**DOI:** 10.1101/829226

**Authors:** Anna Brestovitsky, Daphne Ezer, Sascha Waidmann, Sarah L. Maslen, Martin Balcerowicz, Sandra Cortijo, Varodom Charoensawan, Claudia Martinho, Daniela Rhodes, Claudia Jonak, Philip A Wigge

## Abstract

The decision of whether to grow and proliferate or to restrict growth and develop resilience to stress is a key biological trade-off. In plants, constitutive growth results in increased sensitivity to environmental stress^1,2^. The underlying mechanisms controlling this decision are however not well understood. We used temperature as a cue to discover regulators of this process in plants, as it both enhances growth and development rates within a specific range and is also a stress at extremes. We found that the conserved chromatin-associated protein DEK plays a central role in balancing the response between growth and arrest in Arabidopsis, and it does this via H2A.Z-nucleosomes. DEK target genes show two distinct categories of chromatin architecture based on the distribution of H2A.Z in +1 nucleosome and gene body, and these predict induction or repression by DEK. We show that these chromatin signatures of DEK target genes are conserved in human cells, suggesting that DEK may act through an evolutionarily conserved mechanism to control the balance between growth and arrest in plants and animals.

## Main

Plants are exposed to daily fluctuations in ambient temperature, which influence growth, development and fitness^3^. Being sessile organisms, plants frequently need to opt between growth and stress resilience, making them good systems to analyse how these trade-offs are made.

Recently, it has become clear that the chromatin landscape—i.e. the positioning of the nucleosome containing histone variants such as H2A.Z and H3.3 across the gene body—is correlated to the responsiveness of a gene to environmental variation^3–7^. For instance, hypersensitive environmental genes are enriched in H2A.Z-nucleosomes in their gene bodies, suggesting its involvement in the regulation of environmental response^4,7^. Warm temperature results in H2A.Z-nucleosome eviction and large-scale transcriptional activation in plants^8,9^. Furthermore, H2A.Z is removed from +1 nucleosomes of temperature-induced genes upon temperature increase allowing activation of expression by transcription factors^9^. The molecular mechanism underlying this response likely involves a re-organisation of the chromatin landscape by chromatin remodelling enzymes. In this paper, we demonstrate that DEK-DOMAIN CONTAINING PROTEIN3 (DEK3), the most abundant member of a family of four DEK-domain containing chromatin re-modellers in Arabidopsis^10^, is a link between the chromatin landscape and environmental responsiveness.

The mammalian orthologue DEK is an oncoprotein involved in the development of cancer, inflammation and stem cell biology^11,12^. Because of its key role in cancer, DEK has been intensely studied in animals, and it has been described to have roles as an H3.3 chaperone, co-transcriptional regulator and in splicing^13–16^. Despite these studies, the genome-wide targets directly regulated by DEK are not known^17^, and it is also unclear whether DEK serves as an activator or inhibitor of transcription. In Drosophila and humans DEK can serve as a H3.3 chaperone, and it remodels nucleosomes into more transcriptionally active chromatin through its histone chaperone activity^15^. By contrast, in mammalian cell lines DEK acts as a positive regulator of heterochromatin formation, maintaining the balance between heterochromatin and euchromatin *in vivo*^14^. DEK therefore appears to act as either an activator or a repressor depending on the context.

In Arabidopsis, DEK was identified as a component of the nucleolus by mass spectrometry^16^. DEK3 can change the topology of protein free DNA *in vitro* and cause transcriptional repression of specific loci by increasing nucleosome density^10^. Correct *DEK3* expression is critical for the degree of response of the plants to some stress conditions, including high salinity and heat shock. Plants with constitutively elevated levels of DEK3 are more sensitive to high salinity, whereas plants deficient in *DEK3* are more salt tolerant^10^.

To further understand the global effects of DEK3 on gene expression in plants, we used chromatin immunoprecipitation DNA-sequencing (ChIP-seq) and RNA-seq to investigate how DEK3 occupancy influences the responsiveness of gene expression to temperature. We show that DEK3 affects temperature-dependent biological trade-offs in Arabidopsis, through the up-regulation of stress response genes and by the suppression of thermo-responsive induction of growth and development genes. Furthermore, inherent chromatin landscape features are sufficient to predict whether a gene will be up- or down-regulated by overexpression of DEK3. Additionally, we show for the first time that DEK3 genetically and physically interacts with H2A.Z and modifies H2A.Z-nucleosome distribution within the gene bodies of DEK3 target genes. We suggest a model whereby feedback between the chromatin landscape and chromatin re-modellers can affect trade-off between growth and arrest, and therefore influence developmental plasticity.

## Results

### H2A.Z and DEK interact

Since increasing temperature does not cause H2A.Z-nucleosome loss *in vitro*^9^, we sought to determine if other factors may contribute to this process in plants. To find H2A.Z interacting proteins, we performed H2A.Z affinity purification (from *HTA11::HTA11-FLAG* expressing lines) coupled with mass spectrometry. This approach identified three homologous DEK-domain containing chromatin re-modellers and 6 previously reported H2A.Z interactors^18^ among 263 proteins found in complex with H2A.Z *in vivo* (Supp. Fig. 1a, Supp. Table.1). Two of them, DEK3 and DEK2, have been found among 7 proteins enriched significantly in samples collected at 27°C compared to 17°C (Fig. 1a, Supp. Table.1) suggesting a possible role in temperature pathway regulated through H2A.Z nucleosomes. There are four DEK-domain containing proteins in *Arabidopsis thaliana*, among them, DEK3 shows a strong and abundant expression in all plant organs^10^. Interestingly, DEK3 shows one of the highest fold change when comparing the binding affinities to H2A.Z complexes between 27°C and 17°C (Fig. 1a, Supp. Table 1). We confirmed the interaction of DEK3 with H2A.Z by co-immunoprecipitating H2A.Z-containing nucleosomes and H2A.Z protein (purified from plants), with purified DEK3 (Fig. 1b). This validation set up mimics *in vitro* binding experiments while preserving post-translational modifications of H2A.Z and DEK3. We find that H2A.Z purified from histone extract coimmunoprecipitated with DEK3 in the comparable levels to H2A.Z nucleosomes obtained from nuclear extract, despite reduced levels of other histones as determined by H3 levels (Fig. 1b). This suggests that other protein factors might not be necessary for the interaction between these two proteins.

**Fig. 1.**
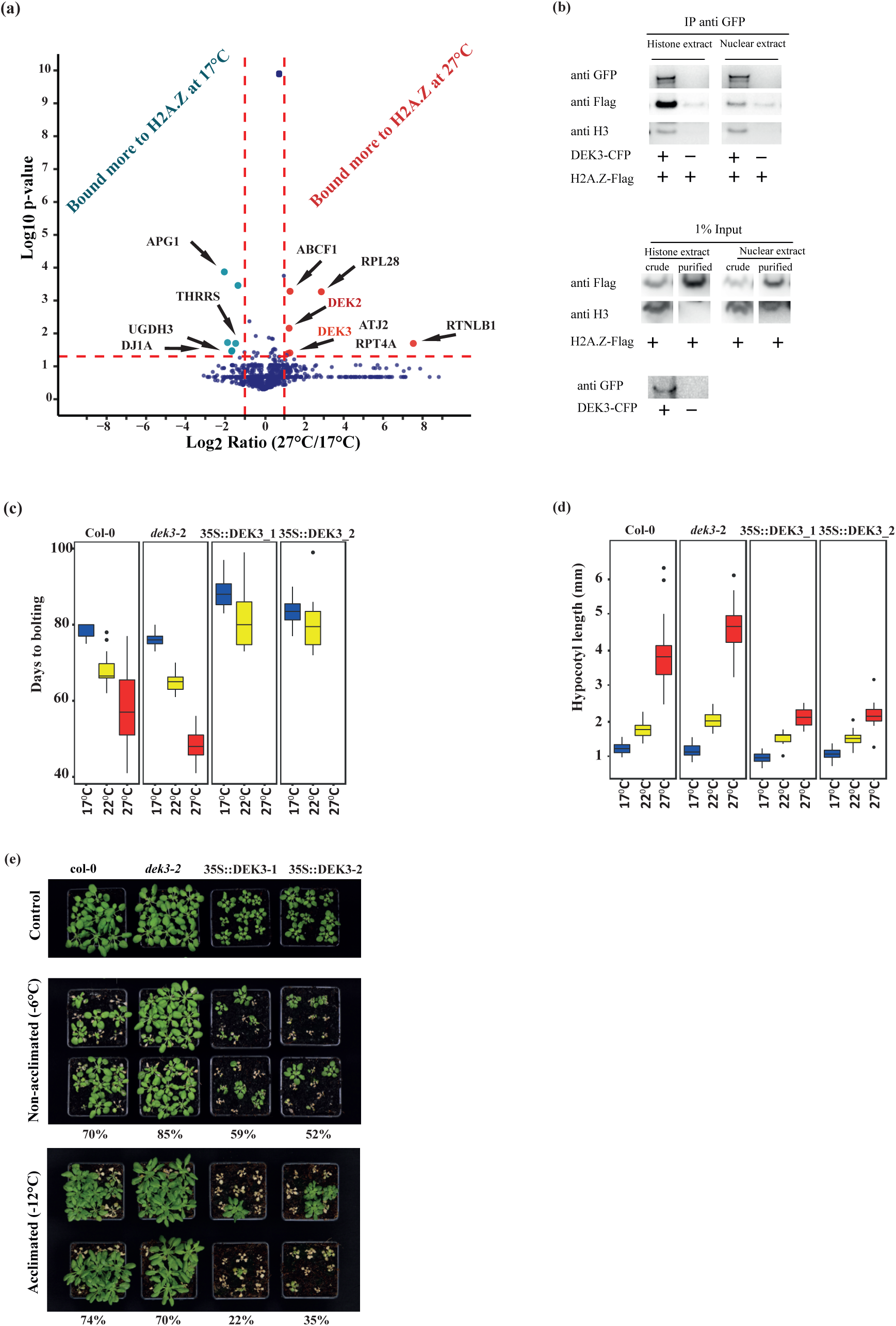
Chromatin protein DEK3 interacts with H2A.Z and plays a role in balancing the temperature response between growth and arrest in Arabidopsis. (a) Volcano plot showing the distribution of proteins identified by Mass Spec in complex with H2A.Z according to *p*-value and fold change binding. Horizontal line indicates significance level at *p*-value ≤ 0.05, and vertical lines shows fold change of 2. (b) Validation of H2A.Z and DEK3 binding. Nuclear extracts from *35S::DEK3-CFP* expressing lines or from Col-0 were subjected to IP with anti-GFP antibodies, and mixed with H2A.Z-Flag purified rom nuclear or histone plant extract. The western blot shows the presence of H2A.Z-Flag, the *35S::DEK3-CFP* and H3 in the immune complexes and in input lysates. The amount of protein in he input represents 1% of the amount of proteins used for IP. (c) Boxplots summarising days to bolting of Col-0, *dek3-2, 35S::DEK3_1* and *35S::DEK3_2* (independent over-expression lines). The plants have been grown at 17°C, 22°C and 27°C. *35S::DEK3_1* and *35S::DEK3_2* plants haven’t bolt within 100 days when grown at 27°C. The ollowing numbers of plants have been used (17/22/27°C): Col-0 9/18/23; *dek3-2* 12/12/24; *35S::DEK3*_1 18/12/24; *35S::DEK3*_2 18/12/24. Box and whisker plots show median, inter-quartile ranges and 95% confidence intervals (t-test p-values are summarized in Supp. Table 5). (d) Boxplots summarising hypocotyl length (mm) of Col-0, *dek3-2, 35S::DEK3_1* and *35S::DEK3_2* (independent over-expression lines). The seedlings have been grown for 7 days under short day photoperiod at 17°C, 22°C and 27°C. The following numbers of seedlings have been used (17/22/27°C): Col-0 55/54/46; *dek3-2* 55/57/58; *35S::DEK3*_1 16/18/16; *35S::DEK3*_2 13/7/17. Box and whisker plots show median, inter-quartile ranges and 95% confidence intervals (t-test statistic p-values are summarized in Supp. Table 5). (e) Summary of the cold sensitivity assays. Col-0, *dek3-2, 35S::DEK3_1* and *35S::DEK3_2* (independent over-expression lines) seedlings have been grown for 21 days in SD at 22°C and either subjected to −6°C for 24hr, or subjected to −12°C for 4 days after 24 h acclimation at 4°C. The plants have been allowed to grow for a further 14 days under recovery conditions, to distinguish between plants that do and do not survive the treatment. All the pictures have been done at the same time and survival rates have been calculated. Percentages indicate survival rate. The survival rate of all control plants has been 100%. The following numbers of plants used in the assay: non-acclimated (survived/total) Col-0 19/27; *dek3-2* 23/27; *35S::DEK3*_1 16/27; *35S::DEK3*_2 14/27; acclimated (survived/total) Col-0 20/27; *dek3-2* 19/27; *35S::DEK3*_1 6/27; *35S::DEK3*_2 7/27.

Previously, immunopurification of DEK3 has identified its interaction with histones H3 and H4, but not with H2A and H2B^10^. The use of an antibody directed against all H2A variants in this previous study made the detection of H2A.Z very difficult as H2A.Z levels are only approximately 10 % of those of H2A^19^.

### Perturbing *DEK* expression levels affects growth and stress response

Temperature has a strong effect on plant growth and development. Growth in high ambient temperatures (below the threshold of inducing widespread heat stress), results in faster growth and accelerated development (thermomorphogenesis), illustrated by increased hypocotyl length and early flowering^20^. In contrast, heat stress induces metabolic imbalance, accumulation of toxic by-products and, adversely influencing reproductive growth and yield quality^21^. Plant resilience to stress conditions such as high salinity and heat shock requires the correct level of DEK3^10^. Additionally, a *DEK3* null allele (*dek3-2*) shows an exaggerated response to warmer (non-stressful) temperature, while overexpression of *DEK3* reduces the thermal responsiveness of seedlings, both as measured by elongation of hypocotyl and flowering time (Fig. 1c-d, Supp. Fig. 1b-c). The altered responsiveness of *DEK3* mis-expressing plants to temperature changes appears to be a general property, since *DEK3* overexpressors are also unable to acclimate to cold temperature and show increased sensitivity to cold stress, applied with and without acclimation (Fig. 1e). *dek3-2* plants are less sensitive to freezing without acclimation (Fig. 1e, middle panel), suggesting a key role of *DEK3* in cold resistance pathways.

We observe similarly altered hypocotyl growth in *dek2* and *dek4* mutants, suggesting a general role for this family of genes in controlling response to temperature (Supp. Fig. 1d).

To further understand why plants with abnormal DEK3 protein levels exhibit altered temperature responses, we investigated gene expression in *dek3-2* and *35S::DEK3* plants at 17 °C and 27 °C. Since the ambient temperature transcriptome is dynamic^22^, we sampled the transcriptome over the 24 h day-night cycle (Supp. Fig. 1e-g). Consistent with plant phenotypes, the largest perturbation in gene expression occurs in *35S::DEK3* at 27 °C (Supp. Fig. 1e). The overexpression of *DEK3* caused ∼4,860 genes to be miss-expressed in at least one time-point compared to WT plants when grown at warm temperature (Supp. table 2).

We used Principal Component Analysis (PCA) to further understand differences in gene expression. Principle Components (PC) 1 and 2 together could explain 54% of the gene expression variance and primarily separate the samples based on time of day into day and night time samples (Supp. Fig. 1f). Among the night samples, the expression of differentially expressed genes in *35S::DEK3* plants at 27 °C resembles the 17 °C transcriptomes of all genotypes rather than that of warm temperature transcriptomes (Supp. Fig. 1f). Indeed, this is consistent with the phenotype of *35S::DEK3* plants, which at 27 °C show perturbed hypocotyl elongation and flowering similar to the plants grown at 17 °C (Fig. 1c-d, Supp. Fig. 1b-c).

Hierarchical clustering of differentially expressed genes reveals three major categories: the first group includes warm temperature induced genes in Col-0, whose expression was induced during the day (first group: clusters 2 and 3) or during the night (Second group: clusters 6 and 7), and genes whose expression was specifically induced only by *DEK3* overexpression at 27 °C (third group: clusters 4 and 5) (Supp. Fig. 1g).

The first group is highly enriched for different GO terms related to metabolism, translation and ribosome biogenesis, RNA methylation, nucleosome assembly and histone modifications (Supp. Table 2); the second group of genes is highly enriched for GO terms connected to shoot and meristem development, regulation of growth, metabolism, transcription (RNA elongation and gene silencing) and photosynthesis (Supp. Table 2). This cluster contains genes encoding heat shock proteins such as *HSP70*, as well as genes related to hypocotyl growth, like *LHY*. Genes in this group are more expressed in *dek3-2* toward the end of the night; Genes in the third group respond only in *35S::DEK3*, and are specifically enriched in GO terms related to response to biotic stimuli and immune response (Supp. Fig. 1g, Supp. Table 2).

### DEK3 direct targets can be activated or repressed by elevated *DEK3* expression

Since the effects of DEK3 on transcription may be indirect, we performed CHIP-seq of *DEK3-CFP* expressed under its native promoter in *dek3-2* mutant plants grown at 17 °C and 27 °C at the end of the night and day. These time points are the most distinct according to a PCA analysis (Supp. Fig. 1f), allowing us to capture the maximal diversity in DEK3 behaviour. We observed DEK3 binding primarily in active chromatin states (Supp. Fig. 2a, the chromatin states 1, 3 and 7) characterized by open chromatin and highly correlated with mRNA-encoding genes^23^.

Since DEK3 primarily binds to DNA in shallow, broad peaks (not sharp peaks like a transcription factor), a peak calling approach was not appropriate for analyzing DEK3 binding profiles. Instead, we identified targeted genes by clustering the DEK3 profiles over the gene bodies. DEK3 can be detected in the majority of gene bodies with defined boundaries at the beginning and the end of the genes (Fig. 2a, Supp. Fig. 2b). For many genes, there is an enrichment for DEK3 in the 3’ end of the gene (Fig. 2a, clusters 5 - 7), which is consistent with its role as an H3.3 chaperone in *Drosophila* and humans^14,15^. Since overexpression of DEK3 protein caused such a dramatic effect on plants physiology and transcriptome at warm temperatures (Fig. 1c-e, Supp. Fig. 1b-c, 1e-g), we checked its chromatin binding profiles and compared them to the profiles of the native DEK3 (Supp. Fig. 2b). Since native and overexpressed DEK3 bind widely throughout the genome (Fig. 2a, Supp. Fig. 2b) it is not possible to make a quantitative comparison in their binding to chromatin, however, it is clear that they have different binding patterns. Overexpression of DEK3 results in extension of binding beyond the ends of the gene body (Supp. Fig. 2b) which might contribute to the altered gene expression seen in *35S::DEK3* (Fig. 1b-d, Supp. Fig.1b-c). We do not observe a significant difference in the binding of DEK3 to chromatin that can be associated with temperature or the time of day (Fig. 2a, Supp. Fig. 2b), suggesting that other factors may be involved in temperature dependent gene expression pattern of DEK3 targets during the day. Hence, we used all four DEK3 ChIP-seq datasets from *DEK3::DEK3-CFP dek3-2* plants as biological replicates for the following analysis in order to determine DEK3 direct targets controlled by temperature.

**Fig. 2.**
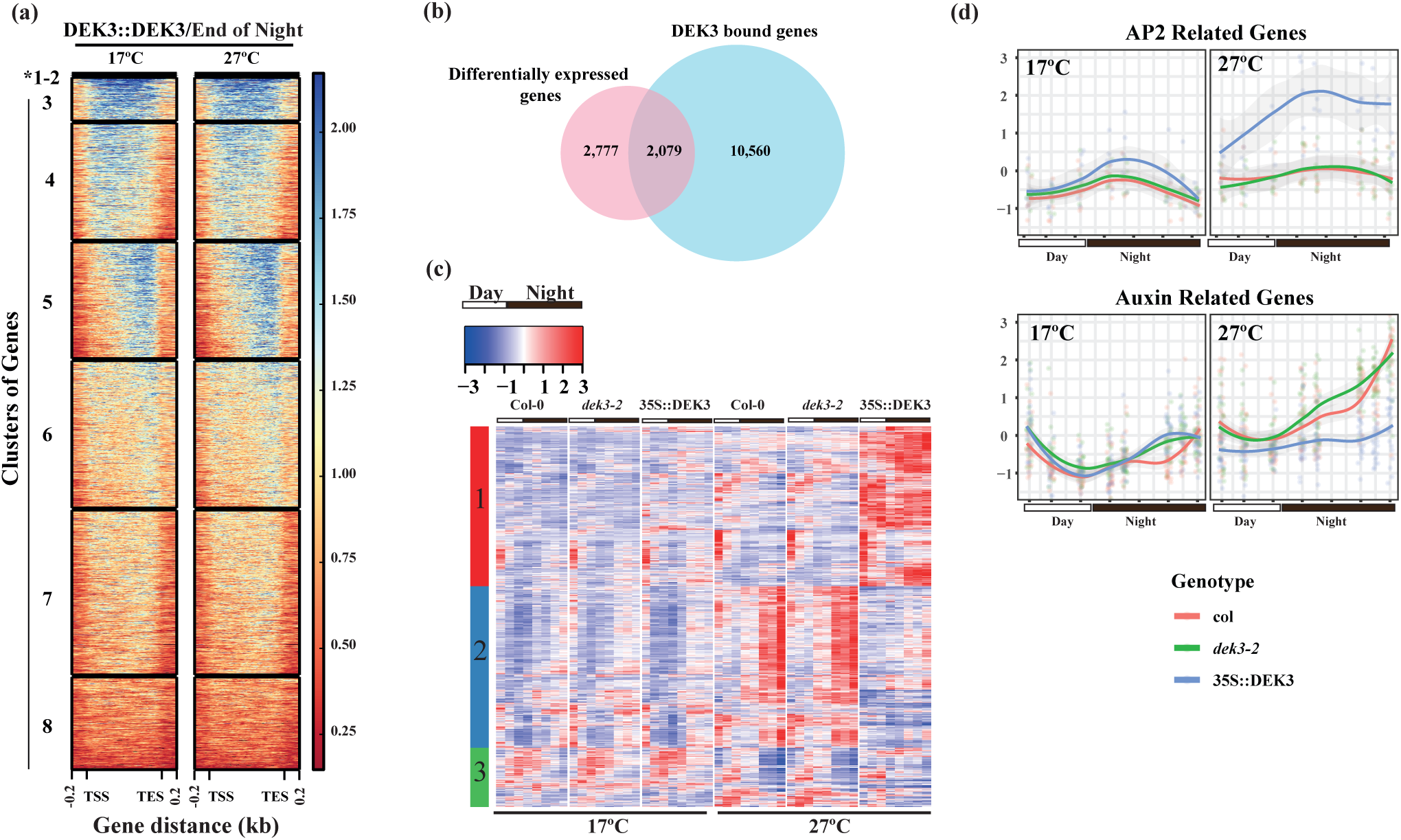
DEK3 directly regulates the expression of “Growth” and “Stress” genes. (a) ChIP-seq binding profiles of DEK3-CFP in seedlings grown at 17°C and 27°C and collected at he end of the night. The DEK3 binding patterns across all gene bodies (including 200 bp upstream of the transcription start site (TSS) and 200 bp downstream of the transcription termination site (TES) have been clustered. A large proportion of genes have an enrichment of DEK3 over their gene body, with clusters 3-5 showing the highest enrichment. The label colour bar on the right shows normalised read count, with the highest level in blue and lower in red. (b) The overlap between the genes mis-expressed by perturbation in DEK3 levels (4,859 genes) and those that are bound directly by DEK3 (12,639 genes, cluster 3-5, from (a)). The overlap highlights DEK3’s direct targets mis-regulated in warm temperature when DEK3 is overexpressed (2,079 genes). (c) Transcriptional patterns of DEK3 direct targets at the end of the night (1,048 genes) which were hierarchically clustered into 3 groups based on their z-scores calculated from transcript per million (TPM) values in all time points. Up-regulated genes are in red and down-regulated genes are in blue. The sidebar to the left of the heatmap indicates the 3 clusters of differentially expressed genes. Black bars on the top indicate night, white bars day. (d) Expression of direct DEK3 targets belong to AP2 and auxin related families as summarised in Supp. Table 2, in Col-0, *dek3-2, 35S::DEK3-CFP* plants grown at 17°C and 27°C under short photoperiod and collected at 8 different time points during 24 hr time course.

We identified 2079 genes as potential direct DEK3 targets (Fig. 2b), as defined by having a high DEK3 occupancy in both temperatures (clusters 3-5 in Fig. 2a) and being differentially expressed at least at one time point at 27 °C when *DEK3* is perturbed (Supp. Table 2).

Since we observe the highest influence of DEK3 levels on gene expression based on PCA and hierarchical cluster analysis at ZT0 (Supp. Fig. 1f-g), we further analysed this timepoint. More than half of the potential direct DEK3 targets (1048 out of the 2079 genes) were differentially expressed at this time point (Supp. Table 3). To analyse the mechanism of DEK3 transcription regulation at this time point, we focused on these 1048 genes for further analysis.

These DEK3 targets can be separated into three groups with distinct transcriptional profiles (Fig. 2c, Supp. Table 3): Genes in Cluster 1 show greatly enhanced expression at 27 °C in *35S::DEK3* compared to Col-0 and *dek3-2* (Fig. 2c). Since these genes are already expressed at low levels in Col-0, it would be challenging to detect a significant reduction in expression in *dek3-2* compared to WT plants (Fig. 2c).

This group is dominated by seven members of the AP2-type transcription factors, mainly of the *ETHYLENE RESPONSE FACTOR (ERF)* class whose expression is induced by DEK3 overexpression in combination with temperature (Fig. 2d). Overall, the genes in this cluster are highly enriched for stress genes (Supp. Fig. 2c). The miss-expression of the stress transcriptome likely contributes to the observed enhanced sensitivity of *35S::DEK3* to freezing (Fig. 1e).

Cluster 2 contains genes induced during the night at 27 °C in Col-0. These genes are up-regulated in *dek3-2* earlier in the night compared to Col-0, while not induced at all when *DEK3* is overexpressed. For example, the temperature responsive auxin biosynthesis gene *YUCCA8* that is necessary for hypocotyl elongation at 27 °C is directly repressed by DEK3. Prominent transcription factors in cluster 2 include the AUXIN RESPONSE FACTORS (ARF) 1 and 19, and the GATA transcription factors 2, 5 and 9. Many genes also implicated in auxin signalling, for example *NAKED PINS IN YUCCA MUTANT (NPY) 1, 3* and *5* are also repressed by DEK3. At the same time these genes are induced even more during the night in *dek3-2* plants in temperature dependent manner (Fig. 2d). Overall there is strong enrichment for the GO-terms associated with growth, development and auxin (Supp. Fig. 2c), indicating that influence on growth caused by *DEK3* perturbations at 27 °C is direct. The final cluster is heterogeneous, but it is the only cluster that contains genes that decrease their expression levels at elevated temperatures.

*DEK3* affects growth and survival in salinity stress^10^, heat stress^10^, elevated ambient temperature and cold stress (Fig. 1c-e, Supp. Fig. 1b-c), indicating that DEK3 controls genes playing a role in promoting stress responses and inhibiting growth in a broad range of conditions. Consistent with this, using publicly available transcriptomic data from the AtGenExpress consortium, we found that the Cluster 1 genes were up-regulated and the Cluster 2 were down-regulated in a wide range of abiotic stress conditions including UV-B, salt, osmotic, cold, wounding, heat, drought, genotoxic and oxidative stresses^24^ (Supp. Fig. 4a).

### The activation or repression of DEK3 targets is related to gene body H2A.Z

*DEK3* mediates the correct expression of a set of genes that are widely responsive to abiotic stresses, while simultaneously repressing the induction of genes promoting growth (Fig. 2c, Supp. Fig. 2c). This raises the question of how a single factor can both activate and repress gene expression in a locus specific manner. This behaviour is similar to that reported in mammals, where DEK behaves as repressor in some cases and as an activator in others^17,25^.

Since chromatin state is correlated with the degree of gene responsiveness^4,6,7^, we investigated whether the binding pattern of H2A.Z, H3.3 and DNA accessibility were predictive of whether a direct DEK3 target would be activated (“stress” genes) or repressed (“growth” genes). To do this, we built a conditional decision tree model that would take the distribution of DEK3, H2A.Z and H3.3 (ChIP-seq) and the DNA accessibility (MNase-seq) in WT as an input, and trained a model to predict whether a DEK3 bound gene would be up- or down-regulated in the *35S::DEK3* plants at 27 °C at the end of the night (Fig. 3a, Supp. Fig. 3a-d), where the biggest difference in expression could be observed (Supp. Fig. 1f).

Strikingly, the most important feature that distinguished between repressed genes (referred to as the “growth” genes) and induced genes (referred to as the “stress” genes) was the gene-body distribution of H2A.Z (p<0.001) with most DEK3 up-regulated “stress” genes having H2A.Z in the gene body, whereas the repressed “growth” genes did not have H2A.Z in the gene body. These “growth” genes predominantly have +1 H2A.Z nucleosomes (Fig. 3a), some “stress” genes were found to have both gene body H2A.Z and strong +1 H2A.Z. H3.3 and chromatin occupancy (via MNase-seq) also contribute to differences in “growth”, and “stress” genes. Specifically, genes with lower H2A.Z levels, but open promoter regions are more likely to be “stress” genes (Fig. 3a, node 2 and genes with lower levels of H3.3 are more likely to be “growth” genes (Fig. 3a, node 6).

**Fig. 3.**
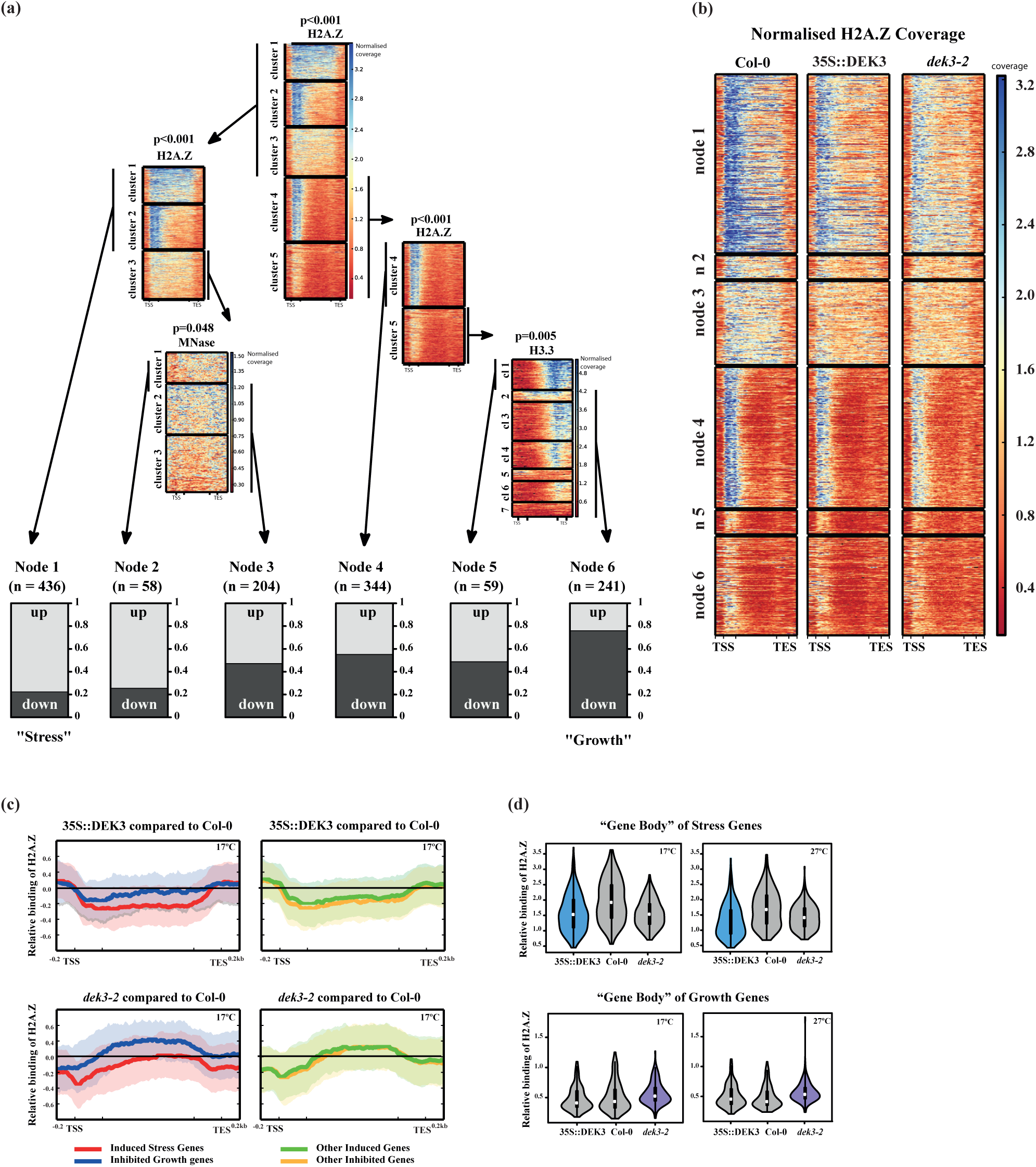
DEK3 affects H2A.Z distribution on target gene bodies. (a) The summary of decision tree predicting whether a DEK3 target gene is up- or down-regulated by overexpression of DEK3 in plants grown at 17°C. Each row in each of the clusters represents a gene, and each column represents a position along the gene body. At each branching point (node), the genes are split into two groups based on their ChIP-seq or MNase-seq profile. (b) The example ChIP-seq binding profiles of H2A.Z-Flag in Col-0, *dek3-2* and *35S::DEK3* plants grown at 27°C and collected at the end of the night. The binding profiles are ordered by the nodes identified by the decision tree. (c) Profiles of H2A.Z-Flag ChIP-seq coverage in *35S::DEK3* and *dek3-2* relative to Col-0 plants, depicted over the gene body, 200bp upstream (TSS) and downstream (TES). Plants were grown at 17°C and collected at the end of the night. (d) The violin plot summarizing the distribution of natively expressed H2A.Z-Flag on the gene bodies of the “stress” and “growth” genes in Col-0, *dek3-2* and *35S::DEK3* plants grown at 17°C and 27°C (Collected at the end of the night).

While the level of DEK3 binding does not determine whether a gene will have H2A.Z in the gene body (Fig. 3a-b, Supp. Fig. 3e), perturbing *DEK3* expression consistently changes the distribution of H2A.Z incorporation around the +1 nucleosome and gene body (Fig. 3c-d, Supp. Fig. 3f). In *dek3-2*, there is a larger ratio of gene body H2A.Z in “growth” genes compared to WT plants (Fig. 3c-d (lower panel), Supp. Fig. 3f (lower panel)), which is consistent with these genes being more easily induced in *dek3-2* compared to Col-0 at 27 °C (Supp. Fig. 4b (upper panel)). This might explain the phenotypic differences between these plants (Fig. 1c-d, Supp. Fig. 1b-c). Conversely, there is lower enrichment for H2A.Z in both the +1 position and gene bodies of “stress” related genes in plants over-expressing *DEK3* compared to WT background (Fig. 3c-d (upper panel), Supp. Fig. 3f (upper panel)).

It has been shown that transcription of genes with H2A.Z in gene bodies is more easily induced by environmental cues and stress^4,7^—these are the genes that are induced by overexpression of *DEK3* and temperature (Fig. 3a, node 1, Supp. Fig. 4b (lower panel)). In contrast, genes that have only +1 H2A.Z nucleosomes and H3.3 around TTS, are expected not to be easily induced by environment but correlated with growth^4^. These genes are suppressed by overexpression of DEK3 and induced in *dek3* in a temperature specific way manner compared to Col-0 (Supp. Fig. 4b (upper panel)).

These observations are consistent with the interaction of DEK3 with H2A.Z (Fig.1a-b, Supp. Fig. 1a), and with decreased expression of the “growth” genes and increased expression of the “stress” related genes in the absence of H2A.Z (Fig. 4d-e) or when its incorporation into chromatin is reduced (Fig. 4f-g). Given that H2A.Z is known to accumulate in the gene body of environmentally responsive genes (such as in drought stress responsive genes)^4,7,6^, a relationship between DEK3 and H2A.Z may suggest a mechanism by which DEK3 perturbations could affect H2A.Z distribution in +1 nucleosomes and gene bodies, and in this way control inducibility of gene expression by environment. This pattern doesn’t hold among strongly DEK3-bound genes in other nodes (Fig. 3c, Supp. Fig. 3f), suggesting that the chromatin landscape is likely to be changing as a result of the DEK3-H2A.Z interaction, rather than as a consequence of changes in transcription.

**Fig. 4.**
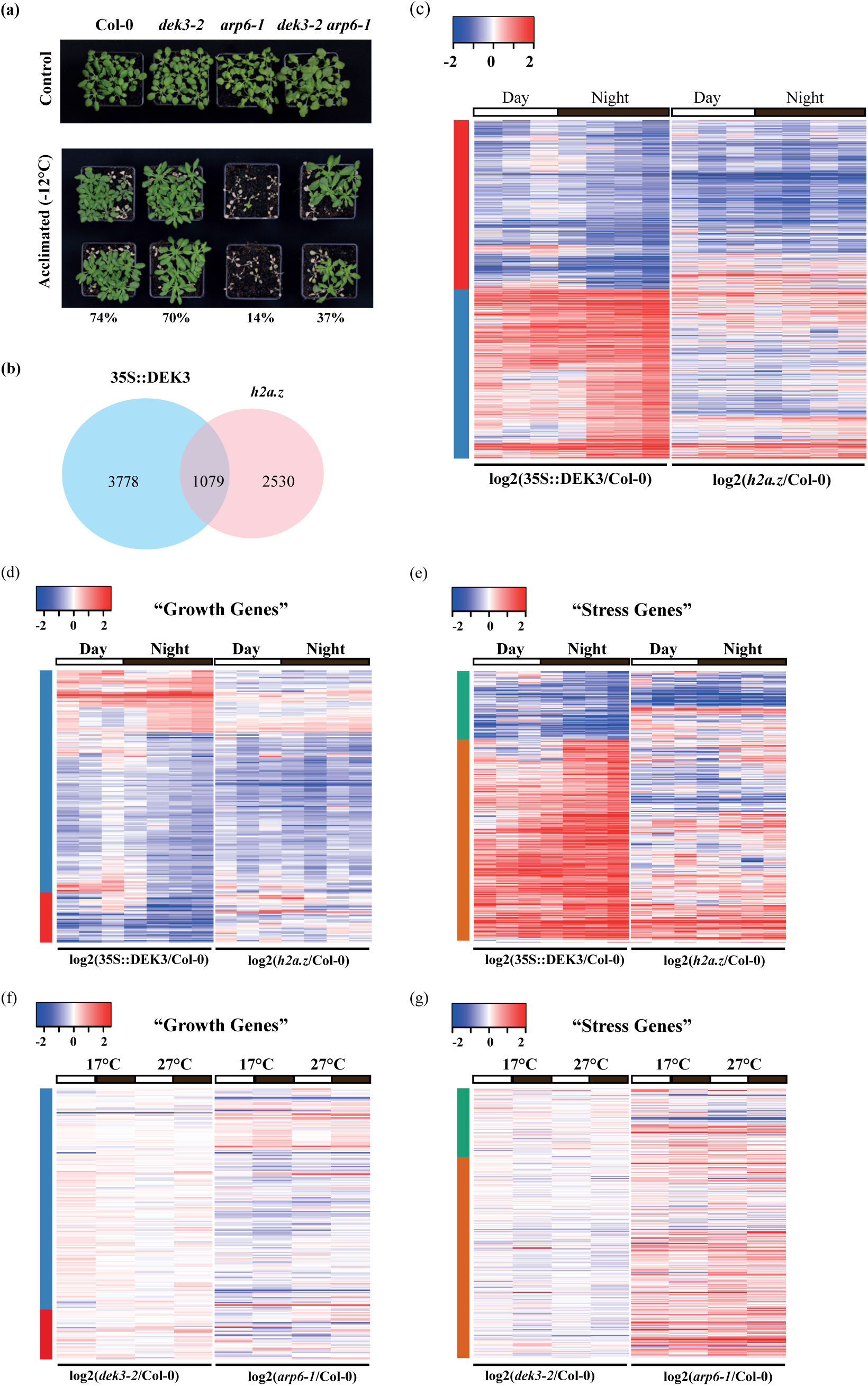
DEK3 controls the expression of “growth” and “stress” genes antagonistically to H2A.Z. (a) Col-0, *dek3-2, arp6-1* and *dek3-2 arp6-1* seedlings were grown for 21 days in SD at 22°C and subjected to −12°C for 4 days after acclimation for 24 hr at 4°C. Percentages indicate survival rate. All control plants have survived. The numbers of plants used: acclimated (survived/total) Col-0 20/27; *dek3-2* 19/27; *arp6-1* 4/28; *dek3-2 arp6-1* 10/27. (b) Venn diagram of overlap between the genes mis-expressed by perturbation in DEK3 levels (4,859 genes) and genes mis-expressed in *h2a.z* plants (3,609). The overlap highlights (1,079 genes) what are mis-regulated in both genotypes in warm temperature. (hypergeometric test p-value of the overlaps < 2.63e-65 for both groups). (c) Expression profiles of DEK3 target genes in *35S::DEK3* and *h2a.z* genetic backgrounds were hierarchically clustered into two groups, based on the log2 ratio compared to the values of Col-0 in ranscript per million (TPM). Plants were grown at 27°C and collected at 8 time points over 24 h. Up-regulated genes are shown in red and down-regulated genes are shown in blue. The sidebar on the left of the heatmap indicates the major clusters. Black bars on the top indicate night, white bars day. (d) (e) Expression of “Growth”(d) and “Stress”(e) genes in *35S::DEK3* and *h2a.z* genetic backgrounds were hierarchically clustered, based on the log2 ratio compared to the values of Col-0 in transcript per million (TPM). Plants has been grown at 27°C and collected at 8 time points during 24 h. Up-regulated genes are shown in red and down-regulated genes are shown in blue. The sidebar on the left of the heatmap indicates major clusters. Black bars on the top indicate night, white bars day. (f) (g) Expression, based on the log2 ratio compared to the values of Col-0 in TPM, of “Growth” (f) and “Stress” (g) genes in *dek3-2* and *arp6-1* genetic backgrounds were hierarchically clustered as in (f) and (g) respectively. Plants were grown at 17°C and 27°C, and collected at the end of the day (white bar) or at the end of the night (black bar). Up-regulated genes are shown in red and down-regulated genes are shown in blue. The sidebar on the left of the heatmap indicates the major clusters.

### *DEK3* and *H2A.Z* interact genetically

Our results suggest that DEK3 plays a role in driving H2A.Z-nucleosome removal from the gene body of stress responsive genes. Supporting this model (Supp. Fig. 4i), when H2A.Z levels or its deposition is perturbed in mutants lacking H2A.Z or *ACTIN RELATED PROTEIN6 (ARP6)*, the transcriptome shows similar mis-expression as in *35S::DEK3* (Fig. 4d-e), but opposite to that of *dek3-2* (Fig. 4f-g). Furthermore, the *arp6-1* mutation is able to partially suppress the effect of *dek3-2* on the transcriptome (Supp. Fig. 4g). Since ARP6 might be present in several protein complexes including SWR1 and may have other roles^26,27^, we also investigated the effect of removing H2A.Z in plants on the *DEK3* dependent transcriptome. In the triple mutant for *hta8,9,11*, we observe a consistent pattern over the time course, with up-regulation of *35S::DEK3* activated genes and down-regulation of *35S::DEK3* suppressed genes (Fig. 4c-e), similarly to what is observed in *arp6-1* (Fig. 4f-g).

In general, “growth” related genes have less H2A.Z signal, and this is most likely to be found near the TSS (Fig. 3a). These temperature dependent genes are further induced by warm ambient temperature in *dek3-2* possibly by increasing the levels of their H2A.Z gene body binding (Fig. 4d, Supp. Fig. 4b (upper panel)). Reducing H2A.Z-nucleosome occupancy in the *h2a.z* and *arp6-1* backgrounds slightly inhibits the temperature induction of “growth” genes (Fig. 4d, 4f). However, when the levels of both proteins, H2A.Z and DEK3, on chromatin are reduced in the *dek3-2 arp6-1* double mutant, the temperature dependent activation of these genes is been impaired (Supp. Fig. 4g (upper panel)). In contrast, the “stress” related genes are enriched in H2A.Z in their gene bodies (Fig. 3a) and up-regulated in the *35S::DEK3* plants grown at 27°C, (Fig. 4e, Supp. Fig. 4b (lower panel)). These genes are also up-regulated in *h2a.z* and *arp6-1* mutants in a temperature-dependent manner (Fig. 4e, 4g), indicating that H2A.Z-nucleosome occupancy may regulate their transcription. This increase is abrogated in *dek3-2 arp6-1* plants, demonstrating the antagonistic interaction between DEK3 and ARP6 (Supp. Fig. 4g (lower panel)).

In light of the physical interaction between DEK3 and H2A.Z (Fig. 1a-b, Sup. Fig. 1a) and in line with the transcriptional patterns of *dek3-2 arp6-1* double mutants and *arp6-1* (Supp. Fig. 4g), we sought to determine if they also interact genetically. The partial rescue of the *arp6-1* phenotype by the *dek3-2* mutation at both an elevated temperature and cold stress (Fig. 4a, Supp. Fig. 4c-f) indicates that *ARP6* and *DEK3* might have opposite functions within the same pathway. Over-expression of *DEK3* in *arp6-1* led to temperature-dependent lethality (Supp. Fig. 4h), suggesting that normal levels of *DEK3* expression are necessary for plants to survive with reduced H2A.Z-nucleosomes when exposed to warm or cold temperature.

### H2A.Z deposition patterns in human DEK targets

DEK proteins are conserved through evolution found in almost all higher eukaryotes^29,30^, and show similarity in domain structure between plants and animals^12^. Three of the four plant DEK proteins, DEK2, DEK3 and DEK4, have been found in complex with H2A.Z by mass spec analysis (Fig. 1a, Supp. Fig. 1a, Supp. Table 1) and showed similar growth phenotypes in response to warm temperature (Fig. 1d, Sup. Fig. 1d), suggesting functional redundancy between them. We therefore asked if our finding that the pattern of H2A.Z-nucleosome occupancy on the gene body may be predictive of how a gene is regulated by DEK3 in Arabidopsis also applies in humans. Previously, human DEK (hDEK) has been shown to both activate and repress expression of the genes involved in cancer and hematopoiesis^30–38^ (summarised in Supp. Fig. 4j). We observe that hDEK up-regulated genes possess mainly gene body H2A.Z, while hDEK down-regulated genes have more +1 nucleosome H2A.Z binding (Supp. Fig. 4j, 4k). This suggests that the interaction between DEK and H2A.Z may be functionally conserved, and that the role of DEK in controlling the balance between growth and stress resilience is conserved between plants and vertebrates. DEK in vertebrates plays key roles in carcinogenesis^7,9,10,13,25,31,36–38,39^, haematopoiesis^25,30,33^ and inflammation^11,46^, processes involving a fine-control of proliferation and stress resilience. It will be interesting to determine if these functions of DEK are also mediated via H2A.Z-nucleosomes.

## Discussion

We find that altered levels of DEK3, an Arabidopsis ortholog of the onco-protein DEK, perturbs developmental programming in *Arabidopsis thaliana* grown in warm temperature by influencing expression of environmental (stress related) and growth related (developmental) genes (Sup. Fig. 2c, Sup. Fig. 4b). The effect of DEK3 on the transcriptome and development appears to be mediated at least in part by changes in the relative distribution of H2A.Z-nucleosomes on the gene bodies of target genes (Fig. 3c-d, Sup. Fig. 3f). These results may explain the dual effect of DEK3 on the transcription of its target genes, shedding light on the longstanding question regarding the influence of DEK on gene expression^12,17,25^. We also provide evidence for physical and genetic interaction between DEK3 and H2A.Z in Arabidopsis (Fig. 1a-b, Sup. Fig. 1a, Sup. Fig. 4c-g). Since DEK proteins are found in most multicellular eukaryotes, as is the H2A variant H2A.Z, these findings may be of broad relevance, and contribute towards understanding of hDEK as a therapeutic target.

The chromatin landscape regulates accessibility of DNA to the transcriptional machinery and transcription factors, controlling the transcriptome and how it responds to a changing environment^47^. Incorporation of the histone variants, H2A.Z and H3.3, into chromatin is frequently associated with responses to environmental perturbations^4,7,8^. While it was reported previously that DEK is a H3.3 chaperone and controls H3.3 deposition into chromatin^14^, the effect of altered levels of DEK3 on more than half of the warm temperature transcriptome can be explained by initial pattern of H2A.Z distribution (Fig. 3a, Sup. Fig. 3f). Previous work suggests that DEK may interact with H3.3^14^ — which is not inconsistent with this study as it is possible for a protein to have multiple binding partners. It will be interesting to see if DEK3 is able to bind H2A.Z and H3.3 simultaneously.

Our results suggest that DEK3 influences the chromatin landscape by modulating the distribution of H2A.Z at specific genes thus controlling their responsiveness to environmental signals (Fig. 3c-d, Sup. Fig. 3f). Our data confirm and extend previous reports suggesting that the initial pattern of H2A.Z on gene bodies is important for the proper induction of transcription in response to environmental stress^4,7^. We show that the levels of H2A.Z on gene bodies is altered by DEK3, influencing gene expression responsiveness in response to environmental signals (Fig. 3). *DEK3* overexpression for example depletes H2A.Z-nucleosome occupancy, leading to enhanced transcriptional responses to higher temperature (Fig. 3c-d, Supp. Fig. 3f, Supp. Fig. 4b).

DEK3 may influence H2A.Z distribution through its role in H3.3 incorporation into chromatin. In animal systems, lack of DEK causes enhanced H3.3 incorporation into the chromatin by HIRA and DAXX/ATRX chaperones^14^. H3.3 incorporation into chromatin might in turn lead to elevated DNA methylation and subsequent prevention of H2A.Z incorporation into gene bodies^6^. This model is not consistent with our finding that increased levels of H2A.Z occur in the gene bodies of *dek3* plants, and conversely, overexpressing *DEK3* results in reduced gene body H2A.Z, at least in the subset of the DEK3 target genes (Fig. 3a, 3c-d, Sup. Fig. 3f). These observations suggest another mechanism of regulation. Interestingly, H3.3 distribution was able to explain the effect of *DEK3* expression on the warm temperature transcriptome for only a subset of the genes (Fig. 3a, node 6), while the H2A.Z pattern alone predicts the behaviour of more than half of the DEK3 target genes (Fig. 3a). This effect could partially be explained by possible physical interaction between DEK3 and H2A.Z (Fig. 1a-b). Even though we could not exclude the possibility of indirect physical interaction between these two proteins, our genetic studies confirm the biological relevance of this interaction (Sup. Fig. 4c-f, Sup. Fig. 4h). Many cancers are associated with impaired DEK levels^12,13,40,41,43,45,46,48–50,14,25,32–35,38,39^, and it will be interesting to determine if these show a similar relationship for transcriptional response and the distribution of hDEK and H2A.Z-nucleosomes.

Our work suggests a model for growth promoting and growth inhibiting responses to environmental changes (Sup. Fig. 4i), where genes with high levels of gene body H2A.Z are responsive to environmental induction, and this sensitivity is enhanced by DEK which can facilitate a decrease in H2A.Z occupancy and higher gene expression. Genes with low gene body H2A.Z occupancy are more likely to be resistant to activation by environmental signals in presence of normal DEK3 levels. For plants this is important for example in tuning the response of the transcriptome to ambient temperature. For plants it is of particular importance to keep a balance between developmental and metabolic transcriptomes under different conditions. Enhanced growth as a result of elevated temperature may cause a reduction in size of adult plant and seed yield^51^. Additionally, plants need to attenuate their development rate in order to provide appropriate response to biotic stress to obtain optimised fitness^2^. However, it may also play a role in other contexts, for example in the case of inflammation and cancer where changes in the relative levels of nutrients and oxygen occur frequently, and cells must respond appropriately.

