## Supplementary material for "DEK influences the trade-off between growth and arrest via H2A.Z-nucleosomes in Arabidopsis": Materials and Methods

### Plant Materials and Growth Conditions

The *dek3-2*, *35S::DEK3-CFP*, *HTA11::HTA11- FLAG* (AT3G54560) and *arp6-1 lines* have been described previously[1–3] . Two independent lines of *35S::DEK3-CFP*, #17-4 and #9-1, have been used there is indicated in the main text, otherwise *35S::DEK3-CFP* #17-4 line has been used[1]. Since HTA9 and HTA11 homologs are very similar in their structure and have redundant functions (March-Diaz et al, 2008), and we have used them in the study interchangeably and collectively called them H2A.Z in the text. Supplementary Table 4 is summarising all the lines and experiments they have been used in.

*35S::HTA9- FLAG* (AT1G52740) line used in part of the biochemical experiments have been constructed by cloning cDNA using Gateway binary pjHA212 vector with Basta resistance containing a C-terminal 3 copies of FLAG tag sequences. The binary construct was transformed into the *hta9-1 hta11-1* double mutant by the floral dipping method. The *35S::HTA9-Flag* transgenic plants were selected on basta-supplemented media and propagated to obtain single insertion lines.

The *h2a.z* mutant has been created by CRISPR CAS9 technology; cloning of an *HTA8*-specific sgRNA sequence (TAGATATGGCTGGTAAAGGTGGG) into the pYB196 vector was performed as described[4]. The construct was introduced into the *hta9-1 hta11-3* double mutant by floral dipping. T1 seedlings were selected for BASTA resistance. T2 plants were then genotyped by PCR-amplifying and sequencing genomic fragments covering the sgRNA target site. A heterozygous plant harbouring a single nucleotide insertion in the first exon of *HTA8*, resulting in a premature stop codon, was identified and propagated to T3 to obtain a homozygous triple mutant.

*dek2* (SALK_137152) and *dek4* (SALK_102885) lines have been received from NASC stock centres, propagated and genotyped to obtain homogenic lines for the following work.

*dek3-2 arp6-1*, *HTA11::HTA11-FLAG dek3-2*, *HTA11::HTA11-FLAG 35S::DEK3-CFP* double mutant lines have been generated by crossing *dek3-2* to the *arp6-1*, *HTA11::HTA11-FLAG* to *dek3-2*, and *HTA11::HTA11-FLAG* to *35S::DEK3-CFP #17-4*, respectively. Their homozygous F3 generations were used for subsequent experiments.

The *DEK3::DEK3-CFP* lines has been generated by amplifying DEK3 together with a 3 kb promoter sequence from genomic DNA and cloned with In-Fusion Advantage PCR cloning kit (#639638; Clontech) into the CFP-containing binary vector pPLV19. The construct was transformed into *dek3-2* by the floral dipping method and transgenic plants were selected on basta-supplemented media and propagated to obtain single insertion lines.

The *HTR5::HTR5-GFP* (At4g40040) for the tagged H3.3 lines has been published previously[5].

*Arabidopsis* seeds were sterilized and sown on ½ x Murashige and Skoog-agar (MS-agar) plates at pH 5.7 without sucrose. Sterilized seeds were stratified for 3 d at 4 °C in the dark and allowed to germinate for 24 h at 22 °C under cool-white fluorescent light at 170 µmol per m2s. The plates were then transferred to short-day conditions (8 h light and 16 h dark) at different temperatures for assays. For hypocotyl length measurement, 7-day-old seedlings were photographed and analyzed using the ImageJ software (http://rsbweb.nih.gov/ij/).

To measure the flowering time the plants have been grown on soil, and we counted the number of days to bolt and/or the number of the leaves on the rosette at the day of bolting.

The hypocotyl length and flowering time measurements have been summarized using boxplots. Each box is bounded by the lower and upper quartiles, the central bar represents the median, and the whiskers are drawn at +/-1.5 x IQR. To test if differences between genotypes have been significant, we performed student t-test. Its results (p values) are summarized in Supp. Table 5.

To assess the sensitivity of the plants to cold, the seedlings have been grown on soil for 21 days in short day conditions at 22°C and either subjected to -6°C for 3 h, or subjected to -12°C for 3 h after 4 days acclimation at 4°C. The temperature was decreased by 2°C/h to reach the respective minimum, and subsequently gradually increased by 2°C/h to reach 4°C after treatment. Plants were then recovered at 22°C for 14 days, to distinguish between plants that do and do not survive the treatment. All the pictures have been done at the same time and survival rates have been calculated. The "survival rate" was defined as a ratio of the number of survived plants to the total number of plants, for each genotype. The survival rate of all control plants in these conditions has been 100%. The numbers of plants used in the assay: non-acclimated (survived/total) Col-0 19/27; dek3-2 23/27; 35S::DEK3_1 16/27; 35S::DEK3_2 14/27; acclimated (survived/total) Col-0 20/27; dek3-2 19/27; 35S::DEK3_1 6/27; 35S::DEK3_2 7/27.

**Immuno-precipitations**

Since there are no commercially available antibodies against Arabidopsis H2A.Z, natively expressed HTA11 was used to pull down endogenous proteins from plant cells. These *HTA11::HTA11-FLAG* lines have been characterised extensively already (Cortijo et al 2017). For immunoprecipitation followed by Mass Spectrometry analysis, *HTA11::HTA11-FLAG* expressing lines or Col-0 seedlings, as control, were grown in liquid ½x MS medium supplemented with 1% sucrose and vitamin B for 7 days at 22 °C in long days and subsequently shifted to 17°C for additional three days. The plants have been shifted to 27°C for 1h or kept at 17°C done1h after dawn on the day of material collection. Plant material was immediately cross-linked when collected using 1% formaldehyde for 15min under vacuum.

Nuclei were purified as previously described (Folta and Kaufman, 2006)[6], resuspended in buffer C [20 mM Hepes–KOH, pH 7.9, 420 mM NaCl, 1.5 mM $MgCl_2$, 0.2 mM EDTA, 0.5 mM DTT, 20% v/v glycerol, 1/10 volume of complete protease inhibitor cocktail (Roche), 1mM MG-132, 0.5mM phenylmethylsulfonyl fluoride (PMSF), 1X phosphotase inhibitor cocktails No2 and No3 (Sigma), 20 mM NaButirate] prepared based on Dignam *et al.* [7]. Nuclear extracts have been sonicated with Bioruptor (Diagenode) and subjected to IP with anti-FLAG antibody conjugated to magnetic beads (SIGMA, cat. No. M8823). The immuno-precipitated complexes have been released by incubation with 100ng/ul 3X FLAG peptide (Sigma, F4799). The released complexes have been boiled and separated on the SDS acryl amide gel (Nupage® 4-12% Bis-Tris Plus Gel, Life Technologies). Gel has been cut into pieces and sent to subsequent Mass Spectrometric analysis.

For validation of Mass Spec results we performed immunoprecipitation experiments followed by western blot. *35S::HTA9-FLAG* and *35S::DEK3-CFP* seedlings have been grown as described above at 22°C for 10 days. Nuclei and nuclear extracts have been obtained as described above, without cross-linking. Histones have been purified by dissolving the nuclei in 0.2 M H2SO4 (0.4N) and precipitation with Trichloroacetic acid (TCA) to denature proteins and release them from binding to other proteins and DNA. The precipitated histones have been finally dissolved in water supplied with 2mM DTT. H2A.Z-FLAG complexes (nuclear extract) and purified H2A.Z protein (histone extract) have been obtained by immuno-purification (IP) with anti-FLAG antibody conjugated to magnetic beads (SIGMA, cat. No. M8823) from nuclear extract and

histone extract respectively; and specifically eluted with 3xFLAG peptide (Sigma, F4799). Nuclear extracts from *35S::DEK3-CFP* expressing lines or from Col-0 have been subjected to IP with anti-GFP antibody (Abcam, cat. NO. ab290) coupled to a 1/1 mix of protein-A and protein-G Dynabeads (life technologies, 10001D and 10003D). The beads have been extensively washed and have been mixed subsequently with H2A.Z-Flag purified from nuclear or histone extract to allow it to bind to DEK3-CFP. Finally, the samples were washed, boiled in SDS loading buffer, and separated by SDS–PAGE. Immunoprecipitated proteins and inputs were analyzed by western blots using specific antibodies. PVDF membranes were developed using Pierce ECL western blotting substrate (32106, Thermo Scientific), and scanned using an Odyssey® Imaging Systems (Li-COR, Biosciences).
The following antibodies have been used in this work: mouse anti FLAG antibody coupled to horseradish peroxidase (HRP) (Sigma, cat. No. A8592); the mouse anti-GFP (Clontech Laboratories, cat, No. 632380) and rabbit anti H3 antibody (Abcam, cat. No. ab 1791). The amount of proteins in the input represents 1% of the amount of proteins used for IP.
The validation has been done in this way to show the possibility of direct interaction between DEK3 and H2A.Z in more physiological way, since we used H2A.Z purified from the most of its original complexes, but retaining its post-translational modifications.

**Mass Spectrometry**

Polyacrylamide gel slices (1-2 mm) containing the purified proteins were prepared for mass spectrometric analysis by manual in situ enzymatic digestion. Briefly, the excised protein gel pieces were placed in a well of a 96-well microtitre plate and destained with 50% v/v acetonitrile and 50 mM ammonium bicarbonate, reduced with 10 mM DTT, and alkylated with 55 mM iodoacetamide. After alkylation, proteins were digested with 6 ng/μL Trypsin (Promega, UK) overnight at 37 °C. The resulting peptides were extracted in 2% v/v formic acid, 2% v/v acetonitrile. The digest was analysed by nano-scale capillary LC-MS/MS using an Ultimate U3000 HPLC (ThermoScientific Dionex, San Jose, USA) to deliver a flow of approximately 300 nL/min. A C18 Acclaim PepMap100 5 μm, 100 μm x 20 mm nanoViper (ThermoScientific

Dionex, San Jose, USA), trapped the peptides prior to separation on a C18 Acclaim PepMap100 3 µm, 75 µm x 250 mm nanoViper (ThermoScientific Dionex, San Jose, USA). Peptides were eluted with a gradient of acetonitrile. The analytical column outlet was directly interfaced via a nano-flow electrospray ionisation source, with a hybrid dual pressure linear ion trap mass spectrometer (Orbitrap Velos, ThermoScientific, San Jose, USA). Data dependent analysis was carried out, using a resolution of 30,000 for the full MS spectrum, followed by ten MS/MS spectra in the linear ion trap. MS spectra were collected over a m/z range of 300–2000. MS/MS scans were collected using threshold energy of 35 for collision-induced dissociation. LC-MS/MS data were then searched against a protein database (UniProt KB) using the Mascot search engine programme (Matrix Science, UK)6. Database search parameters were set with a precursor tolerance of 5 ppm and a fragment ion mass tolerance of 0.8 Da. Two missed enzyme cleavages were allowed and variable modifications for oxidized methionine, carbamidomethyl cysteine, pyroglutamic acid, phosphorylated serine, threonine and tyrosine were included. MS/MS data were validated using the Scaffold programme (Proteome Software Inc., USA). All data were additionally interrogated manually.

Results of tree replicates have been combined using Scaffold programme. EmPAI values have been calculated and data have been exported as "Samples Report Summary". We used one-tail paired t-test to identify significantly enriched proteins in "treated samples", either 17°C or 27°C, relative to "Control samples". In this way the "batch effect" was reduced. Proteins with more than 2-fold change binding compared to control and with $p$-value ≤ 0.1 have been considered to be specifically purified with H2A.Z (Supp. Table 1). 6 previously reported H2A.Z interactors[8] have been detected (Supp. Table 1). One-tail paired t-test was used to identify proteins enriched in one temperature compared to other. Proteins with more than 2-fold change between temperatures and with $p$-value ≤ 0.05 have been considered to be specifically enriched (Supp. Table 1). Volcano plot was created using statistical program R (Fig. 1a).

**RNA-Seq Library Preparation**

The following plants were used for time-course RNA-seq experiments: The Col-0, *dek3-2* and *35S::DEK3-CFP* #17-1 seedlings were grown for 7 days at 17 °C and 27 °C. The plants were sampled at intervals over the diurnal cycle: ZT = 0, 1, 4, 8, 12, 18, 20 and 24 h. Two biological replicates have been sampled; The Col-0, *arp6-1, dek3-2* and *dek3-2 arp6-1* double mutants have been grown at 17°C and 27°C and collected at the end of the day (ZT=8) and at the end of the night (ZT=0). Two biological replicates have been sampled; The Col-0 and *h2a.z* samples have been grown at 27°C and sampled at: ZT = 0, 1, 4, 8, 12, 16, 20 and 24 h. One biological replicate has been sampled.

Total RNA was isolated from 30 mg of ground seedlings using the MagMAX-96 Total RNA Isolation kit (Ambion, AM1830), following the manufacturer's instructions. RNA quality and integrity were assessed on the Agilent 2200 TapeStation. Library preparation was performed using 1 mg of high-integrity total RNA (RIN>8) using the NEBNext® Ultra™ Directional RNA Library Prep Kit for Illumina (NEB, cat. No. NEB #E7420). Libraries were sequenced in-house on an Illumina NextGen 500 sequencer using paired-end sequencing of 75 bp in length.

**RNA-Seq Mapping and Differential Expression Analysis**

The raw reads obtained from the sequencing were analyzed using a combination of publicly available software and in-house scripts as described in Cortijo et. al.[3] No mis-match was allowed when mapping using TopHat.

The genes whose transcription affected by DEK3 expression were identified using DESeq[9] through R Bioconductor. Further analyses on these genes were performed using their TPM values[10], whereby different number of reads in libraries and transcript lengths were taken into account and normalized. The genes with a Padj<1 and TPM >1 were considered to be differentially expressed and were used for further analysis.

Differentially expressed genes in *h2a.z* mutant have been defined base on Log2(TPM sample /TPM Col-0) being greater than 1.5, after filtering out genes whose expression was lower than 1 and genes with low expression variation (coefficient of variation = standard deviation of expression/mean expression < 0.2). One biological replicate has been analyzed. Since temporal pattern of expression of DEK3 target genes didn’t change in *h2a.z* background (Fig. 4c-e), we considered each time point as a separate replicate for the following analysis.

Hierarchical clustering of transcriptomic data was performed using the statistical program R, using the function hclust on 1-Pearson correlation. z-scores have been calculated for each gene based on their mean TPM values from two replicates across all the time points and all the genotypes. In order to create all the heatmaps, except for the Fig.4 (c-g), we have used z-scores. Log2(TPM sample /TPM Col-0) for each time point have been used to generate Fig.4 (c-g).

The dynamic patterns of expression changes in different gene clusters were represented using the heatmap.2 function in R.

Hypergeometric test has been used to find the significance of the overlaps in vein diagrams.

All raw reads were deposited on Gene Expression Omnibus (GSE114646,) or on SRA (SUB4039352, SUB2445017).

To assess over-represented biological functions of the DEK3 target genes in different clusters, we performed the GO enrichment analysis using the Gene Ontology enrichment analysis and visualisation R package "GOsummaries"[11].

Transcriptomic data used in Supp. Fig. 4a have been downloaded from AtGenExpress consortium[12] .

**Chromatin Immunoprecipitation (ChIP)**

The following plants were used for CHIP-seq experiments:

The *HTA11::HTA11- FLAG*, *HTA11::HTA11-FLAG dek3*-2, *HTA11::HTA11-FLAG 35S::DEK3-CFP* double mutant lines has been used to follow after H2A.Z binding profiles in different DEK3 expressing lines; *DEK3::DEK3-CFP dek3-2* has been used to identify DEK3 target genes; *35S::DEK3-CFP* #17-4[1] CHIP-seq and previous published data[1] have been used to test DEK3 binding profiles when this protein is over-expressed; *H3.3-GFP* [5] CHIP-seq and published previous data[5] have been used to check the H3.3 binding profiles to DEK3 direct targets influenced by temperature.

The seeds were germinated at 22 °C and seedlings were grown for 9 days at 17 °C and 27 °C. The plants were sampled at the end of night and day (ZT = 0 and ZT=8) as indicated in the text. Two biological replicates have been sampled for CHIP-seq experiments, except for the *H3.3-GFP* and *DEK3::DEK3-CFP* ChIP-seq, which have been done in one replica for several conditions indicated in main text.

*35S::DEK3-CFP* and *H3.3-GFP* ChIP samples have been collected after plants have been shifted 1 hr after dawn (ZT=1) from 17°C to 27°C for 1hr. Additional replica of *H3.3-GFP* ChIP data has been described previously (Wollmann et al.)[5].

The following plants were used for MNAse-seq experiments:

The *HTA11::HTA11-FLAG*, *HTA11::HTA11-FLAG dek3-2*, *HTA11::HTA11-FLAG 35S::DEK3-CFP* double mutant lines.

Preparation the ChIP experiment was performed as described by Gendrel et al. (2002)[13] with minor modifications. Plant material was immediately cross-linked when collected using 1% formaldehyde for 15min under vacuum. Chromatin was extracted from 1 g of cross-linked materials. For MNAse-seq and ChIP -seq of: *HTA11::HTA11- FLAG*, *HTA11::HTA11-FLAG dek3-2*, *HTA11::HTA11-FLAG 35S::DEK3-CFP* double mutant lines and H3.3-GFP, the chromatin was resuspended in MNase digestion buffer (20mM Tris-HCl [pH8], 50mM NaCl, 1mM DTT, 0.5% NP-40, 1mM CaCl2, 0.5mM phenylmethylsulfonyl fluoride (PMSF) and 1X protease inhibitor cocktail [Roche]), and digested with 0.4U/ml of micrococcal nuclease (MNase, Sigma, N3755) for 15 min. Digestion was stopped with 5 mM EDTA. MNase digestion was performed to obtain a mono-nucleosome resolution as MNase preferentially digests nucleosome-free DNA and the linker regions, whereas sequences bound by nucleosomes are relatively protected from the digestion[14]. For MNAseseq samples have been used for library preparation directly after digestion. For DEK3::DEK3-CFP and 35S::DEK3-CFP ChIP-seq, chromatin was fragmented by sonication using a Bioruptor (Diagenode) in lysis buffer (10 mM Tris-HCl [pH 8], 150 mM NaCl, 1 mM EDTA [pH 8], 0.1%

deoxycholate, and 1X protease inhibitor cocktail). All ChIPs were performed in a buffer containing 1 20mM Tris-HCl (pH8), 150mM NaCl, 2mM EDTA, 1% triton X-100 and 1X protease inhibitor cocktail. ChIP were performed for *HTA11::HTA11-FLAG* using FLAG M2 magnetic beads (Sigma, M8823) and for *DEK3::DEK3-CFP*, *35S::DEK3-CFP*, H3.3-GFP using anti GFP antibody (Abcam, ab290) coupled to a 1/1 mix of protein-A and protein-G Dynabeads (life technologies, 10001D and 10003D). All samples have been subsequently washed 2 times with low salt buffer (20mM Tris buffer (pH=8), 150mM NaCl, 0.1%SDS, 1% Triton) twice with high salt buffer (20mM Tris buffer (pH=8), 500mM NaCl, 0.1%SDS, 1% Triton), once with LiCl buffer (10mM Tris buffer (pH=8), 250mM LiCl, 1%Sodium deoxycholate, 1% NP40) and twice with TE buffer (10mM Tris buffer (pH=8), 1mM EDTA). The elution was done for anti-Flag ChIP with 100ng/ul of 3XFLAG in TE buffer and for anti-GFP ChIP with elution buffer (1% SDS, 0.1M NaHCO3). DNA was extracted using AMPure beads using a ratio of 2.3X as described previously[15].

MNAseseq and ChIP-seq libraries have been prepared in house using the NEBNext® Ultra™ II DNA Library Prep Kit for Illumina® (#E7645). The libraries were sequenced using paired-end 75bp on NextSeq500 on site.

**Analyses of ChIP-Seq and Nucleosome Profiles and Building Decision Tree**

Sequenced ChIP-seq data were analyzed in house, following the same quality control and pre-processing as in RNA-seq. The read counts mapped to each base pair in each sample were normalized by the sample's genome-wide mappable reads coverage per base pair, and used in the subsequent statistical analyses.

To characterise chromatin features associated with DEK3 chromatin binding, we used previously published landscape of *Arabidopsis thaliana* chromatin, which has been divided into 9 states based on the different epigenetic marks[16].

The majority of genes do not change their expression between 17ºC and 27ºC, suggesting that many of the chromatin landscapes are indeed maintained in different temperatures[3,17]. This allowed us to use the previously published landscape of *Arabidopsis thaliana* chromatin[16] to characterise

chromatin features associated with DEK3 chromatin binding. The accumulation of DEK3 as found by ChIP-seq has been divided into 9 states based on the different epigenetic marks[16] and plotted using box-plot. The chromatin states 1, 3 and 7 characterize open chromatin and are highly correlated with gene expression; States 2, 4, 5 and 9 contain the lowest amount of mRNA-encoding genes; Chromatin state 8 is correlated with GC-enriched heterochromatin containing transposable elements [16].
After qualitatively observing the DEK3-ChIP-seq data as a necessary first step in data analysis, we noticed that DEK3-ChIP-seq did not have the structure of small regions with sharp peaks of binding, but rather show shallow enrichment over long stretches, in particular over gene bodies. Peak calling identifiers, like MACS are looking for small regions with sharp peaks of binding (i.e. steep slopes) and not applicable in this case. Chromatin remodelers and histones are widely bound to chromatin; therefore we used their binding profiles, rather then peak calling to determine direct targets of DEK3 as well to characterise histone binding and nucleosome profiles.

ChIP-seq and MNase-seq profiles over gene bodies were drawn using DeepTools[18]. The TSS and TES for each gene were identified using TAIR10, and a 200bp unscaled region was included before and after the gene body, and in addition, the first 300bp of the gene from the TSS were unscaled in order to allow the user to observe the first two nucleosome positions at the start of the gene, which are often well-positioned. K-means was used for clustering.
To investigate the epigenetic basis for DEK3 dual regulation of the transcription, we investigated the role of H2A.Z, H3.3 and DNA acceptability in the determination of whether a predicted target gene would be up- or down-regulated in 35::DEK3-CFP at 27°C. To do this, we built a conditional decision tree model that would take as input the distribution of DEK3, H2A.Z, and H3.3 (ChIP-seq) and the DNA accessibility (MNase-seq) in Col-0 (see Fig. S4) and train a model to predict whether a DEK3 bound gene would be up- or down- regulated in 35::DEK-CFP plants at 27°C. The input into the decision tree model was the clusters of the ChIP-seq and MNase-seq data over the gene bodies of direct targets, using k-means in DeepTools[18], and the

output was whether the gene was up- or down- regulated. Since we had H2A.Z ChIP-seq data available at the end of the night (ZT0), we only used genes that were differentially expressed in 35::DEK3-CFP at the end of the night in this analysis. A conditional decision tree (calculated using the “party” package in R[19]) recursively tests whether the null hypothesis (that the input variables and the output variables are independent). If the null hypothesis cannot be excluded, the recursive algorithm stops. Otherwise, a new branch in the tree is added which splits the tree on the independent variable that is most associated with the output variable.

All raw reads were deposited on Gene Expression Omnibus (GSE114185).

The profiles of human H2A.Z have been created based on the previous published data[16].

1. Waidmann, S., Kusenda, B., Mayerhofer, J., Mechtler, K. & Jonak, C. A DEK Domain-Containing Protein Modulates Chromatin Structure and Function in Arabidopsis. *Plant Cell Online* **26,** 4328–4344 (2014).
2. Deal, R. B. The Nuclear Actin-Related Protein ARP6 Is a Pleiotropic Developmental Regulator Required for the Maintenance of FLOWERING LOCUS C Expression and Repression of Flowering in Arabidopsis. *Plant Cell Online* **17,** 2633–2646 (2005).
3. Cortijo, S. *et al.* Transcriptional regulation of the ambient temperature response by H2A.Z-nucleosomes and HSF1 transcription factors in Arabidopsis. *Mol. Plant* **10,** 1258–1273 (2017).
4. Materials_and_methods_Brestovitsky_v2_MB230418.
5. Wollmann, H. *et al.* Dynamic deposition of histone variant H3.3 accompanies developmental remodeling of the Arabidopsis transcriptome. *PLoS Genet.* **8,** e1002658 (2012).
6. Folta, K. M. & Kaufman, L. S. Isolation of Arabidopsis nuclei and measurement of gene transcription rates using nuclear run-on assays. *Nat. Protoc.* **1,** 3094–3100 (2007).
7. Dignani, J. D., Lebovitz, R. M. & Roeder, R. G. Accurate transcription initiation by RNA polymerase II in a soluble extract from isolated mammalian nuclei. *Nucleic Acids Res.* **11,** 1475–1489 (1983).
8. March-Díaz, R. & Reyes, J. C. The beauty of being a variant: H2A.Z and the SWR1 complex in plants. *Mol. Plant* **2,** 565–77 (2009).
9. Simon, W. Differential expression analysis for sequence count data. *Genome Biol.* **11,** (2010).
10. Wagner, G. P., Kin, K. & Lynch, V. J. Measurement of mRNA abundance using RNA-seq data: RPKM measure is inconsistent among samples. *Theory Biosci.* **131,** 281–285 (2012).
11. Kolde, R. & Vilo, J. GOsummaries: an R Package for Visual Functional Annotation of Experimental Data. *F1000Research* 1–23 (2015). doi:10.12688/f1000research.6925.1
12. Kilian, J. *et al.* The AtGenExpress global stress expression data set: Protocols, evaluation and model data analysis of UV-B light, drought and cold stress responses. *Plant J.* **50,** 347–363 (2007).
