## Supplementary figures for "DEK influences the trade-off between growth and arrest via H2A.Z-nucleosomes in Arabidopsis"

#### Supplementary Fig. 1.

#### DEK3 controls the plant temperature response and transcriptome

(a) The summary of enrichment of DEK family proteins found in complexes immunopurified with H2A.Z-FLAG as has been detected by Mass Spec analysis in 3 independent experiments.

(b) Representative photo showing differences in hypocotyl growth of Col-0, *dek3-2*, *35S::DEK3_1* and *35S::DEK3_2* seedlings grown for 7 days under short day photoperiod at 17°C, 22°C and 27°C. Scale bars, 10mm.

(c) Representative photo showing difference in flowering of Col-0, *dek3-2*, *35S::DEK3_1* plants grown under short day photoperiod at 17°C and 27°C. *35S::DEK3_1* plants have not flowered within 100 days when grown at 27°C.

(d) Boxplots summarising hypocotyl length (mm) of Col-0, *dek2*, *dek4*. The seedlings have been grown for 7 days under short day photoperiod at 17°C, 22°C and 27°C. The following numbers of seedlings have been used (17/22/27°C): Col-0 20/19/18; *dek2* 22/21/19; *dek4* 22/21/23. Box and whisker plots show median, inter-quartile ranges and 95% confidence intervals.

(e) Clustering of all the samples of RNA-seq time courses for Col-0, *dek3-2*, *35S::DEK3* plants grown at 17°C and 27°C under short photoperiod and collected at 8 different time points during 24 hr time course.

(f) Principal component analysis (PCA) of transcriptomes of Col-0, *dek3-2*, *35S::DEK3* plants grown at 17°C and 27°C under short photoperiod and collected at 8 different time points during 24 hr time course. Genes have been used as features. Analysis have been performed on the differentially expressed dataset in *35S::DEK3* plants compared to Col-0 WT grown at 27°C as determined by DEseq[1].

(g) Transcriptional patterns and dynamics of differentially expressed genes (4,859 genes) in *35S::DEK3* compared to Col-0 WT plants grown at 27°C and collected at 8 time points over 24 h. The differentially expressed genes were hierarchically clustered into 8 groups, based on the z-scores calculated from transcript per million (TPM) values at all time points. Up-regulated genes are in red and down-regulated genes are in blue. The sidebar to the left of the heatmap indicates the eight clusters of differentially expressed genes. Black bars on the top indicate night, white bars indicate day.

### Supp. Figure 1 (page 1)

(a)

| Experiment | | | Quantitative value (emPAI) 17C | 27C | Control | 27C/17C |
|---|---|---|---|---|---|---|
| #1 | **DEK2** | AT5G63550 | 0.11 | 0.23 | 0.06 | 2.1 |
| | **DEK3** | AT4G26630 | 0.69 | 1.20 | 0.04 | 1.7 |
| | **DEK4** | AT5G55660 | 0.19 | 0.69 | 0.00 | 3.6 |
| #2 | **DEK2** | AT5G63550 | 0.11 | 0.30 | 0.11 | 2.6 |
| | **DEK3** | AT4G26630 | 0.35 | 0.90 | 0.00 | 2.6 |
| | **DEK4** | AT5G55660 | 0.04 | 0.15 | 0.00 | 4.0 |
| #3 | **DEK2** | AT5G63550 | 0.11 | 0.28 | 0.00 | 2.5 |
| | **DEK3** | AT4G26630 | 0.08 | 0.27 | 0.08 | 3.5 |
| | **DEK4** | AT5G55660 | 0.20 | 0.04 | 0.07 | 0.2 |

(b)

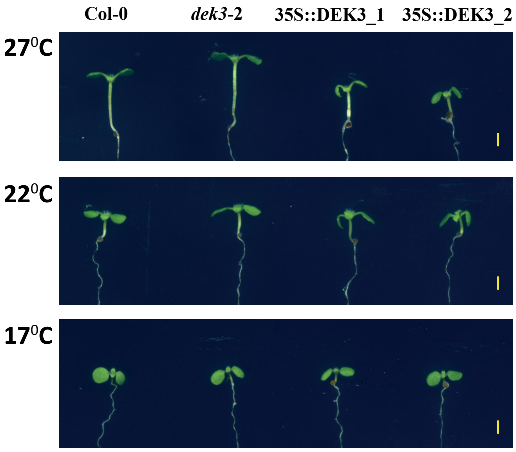


(c)

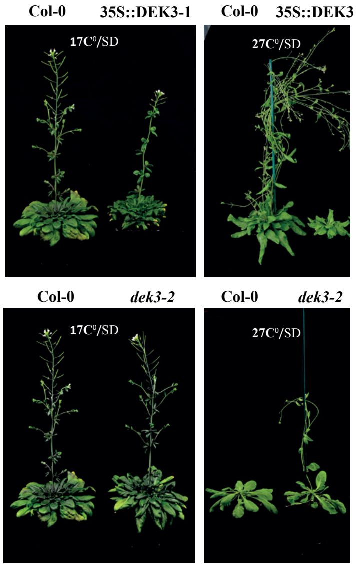


(d)

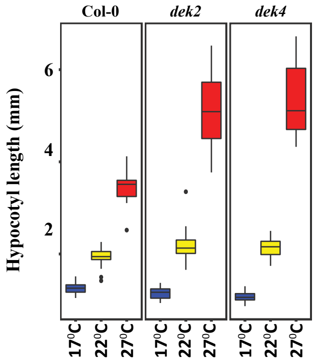

### Supp. Figure 1 (page 2)

(e)

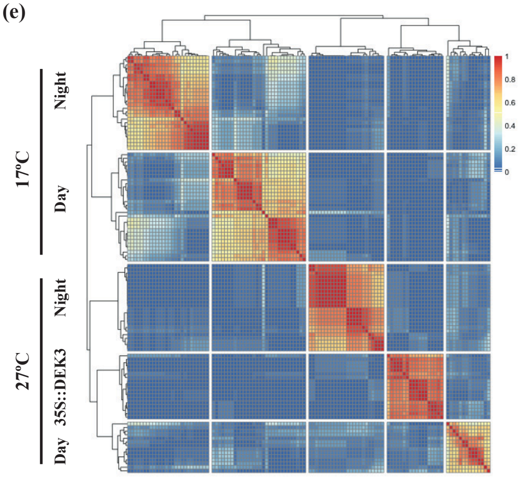


(f)

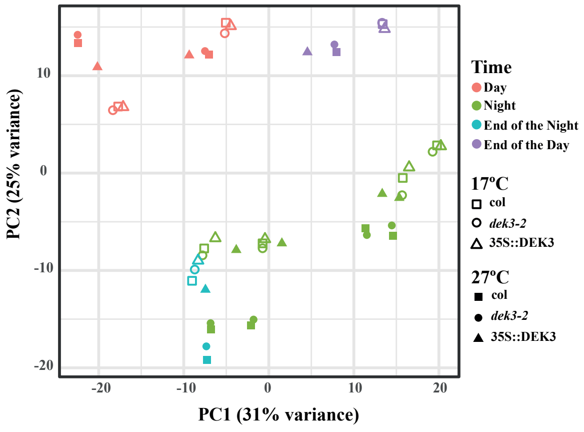


(g)

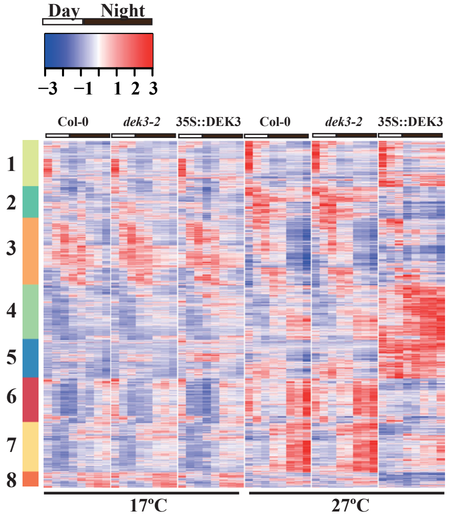

#### Supplementary Fig. 2.

#### DEK3 directly regulates the expression of “Growth” and “Stress” genes.

(a) Boxplot shows the natural log of the ratio between the observed and expected DEK3 read counts over each chromatin state defined by the 9-state chromatin model[2] which is based on clustering genomics regions by their epigenomic signatures. The chromatin states are colour-coded by their functional annotations. Distribution of DEK3 by chromatin state shows that DEK3 binds in gene-rich and transposon-rich chromatin states. The chromatin states 1, 3 and 7 characterize open chromatin and are highly correlated with gene expression; States 2, 4, 5 and 9 contain the lowest amount of mRNA-encoding genes; Chromatin state 8 is correlated with GC-enriched heterochromatin containing transposable elements[2].

(b) ChIP-seq binding profiles of natively expressed DEK3-CFP (*DEK3::DEK3*) and over-expressed DEK3-CFP (*35S::DEK3*) in seedlings grown at 17°C and 27°C and collected at the end of night or end of day. The ChIP binding patterns across all gene bodies (and the 200bp upstream (TSS) and downstream regions (TES)) were clustered. Overexpressed DEK3 (*35S::DEK3*) shows more diffuse binding with less clear borders in the beginning (TSS) and the end (TES) of the genes. The label colour bar on the right depicts normalised read count, with the highest level in blue and lower in red.

(c) Over-represented biological functions of DEK3 direct targets in cluster 1 and cluster 2 (Fig. 2c) summarized by world clouds. A violin plot shows gene expression, and a word cloud showing the most significant GO results describes each cluster. The sizes and colours of the taxons in word clouds are proportional to - $log_{10}$ of the Spearman rank correlation test p-value. The absolute enrichment strength of terms (words) is color-coded in grey scale. The violin plot summarises the relative expression data for each cluster in *35S::DEK3* and Col-0 plants grown at 27ºC and collected over 24 h. Black bars on the bottom indicate night, white bars day.

### Figure 2 supp

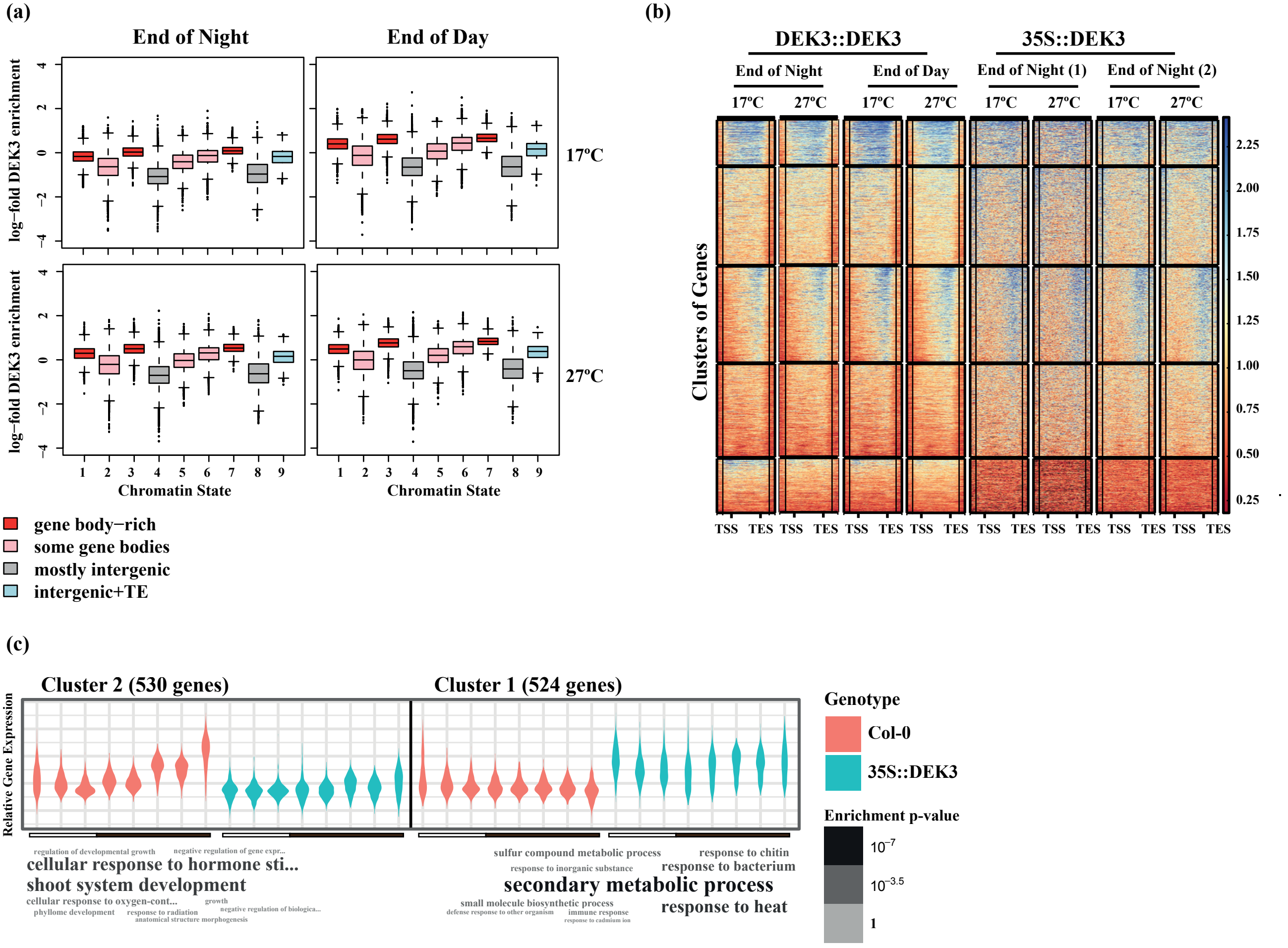

#### Supplementary Fig. 3.

#### Construction of a Decision Tree and influence of DEK3 levels on the H2A.Z distribution on the gene bodies

(a) Construction of a Decision Tree: the direct targets are defined as genes that are DEK3-bound and that are differentially expressed in *35S::DEK3* plants compared to wild type. Among these, a proportion of the genes were defined as “Stress” genes and “Growth” genes.

(b) The goal of the decision tree was to utilize data from MNase-seq, H2A.Z, H3.3, and DEK3 ChIP-seq experiments to predict whether a gene should be classified as a 'stress' gene or a 'growth' gene. To do this, a tree was constructed in which at each branching point a yes-or-no question is asked, and the answer to the question informs whether the gene is more likely to be a 'stress' gene or a 'growth' gene.

(c) This schematic shows the results depicted in Fig. 3a and describes what these 'questions' are in words.

(d) The summary showing the input values into the decision tree model (clusters of MNase-seq and ChIP-seq data). The decision tree gave an output of 6 different nodes, and the bottom panel shows the MNase-seq and ChIP-seq data arranged by node.

(e) Profiles of H2A.Z-Flag ChIP-seq coverage in Col-0, *dek3-2* and *35S::DEK3* plants depicted over the gene body (and 200bp upstream (TSS) and downstream (TES)). This shows all the H2A.Z ChIP-seq data in all conditions tested in two replicates, arranged by the nodes output from the decision tree. The label colour bar on the right is showing normalised read count, with the highest level in blue and lower in red. Plants has been grown at 17°C and 27°C, and collected at the end of the night.

(f) Relative profiles of H2A.Z-Flag ChIP-seq coverage in *35S::DEK3* and *dek3-2* relative to Col-0 plants, depicted over the gene body, 200bp upstream (TSS) and downstream (TES)). Plants were grown at 27°C and collected at the end of the night.

### Figure 3 supp

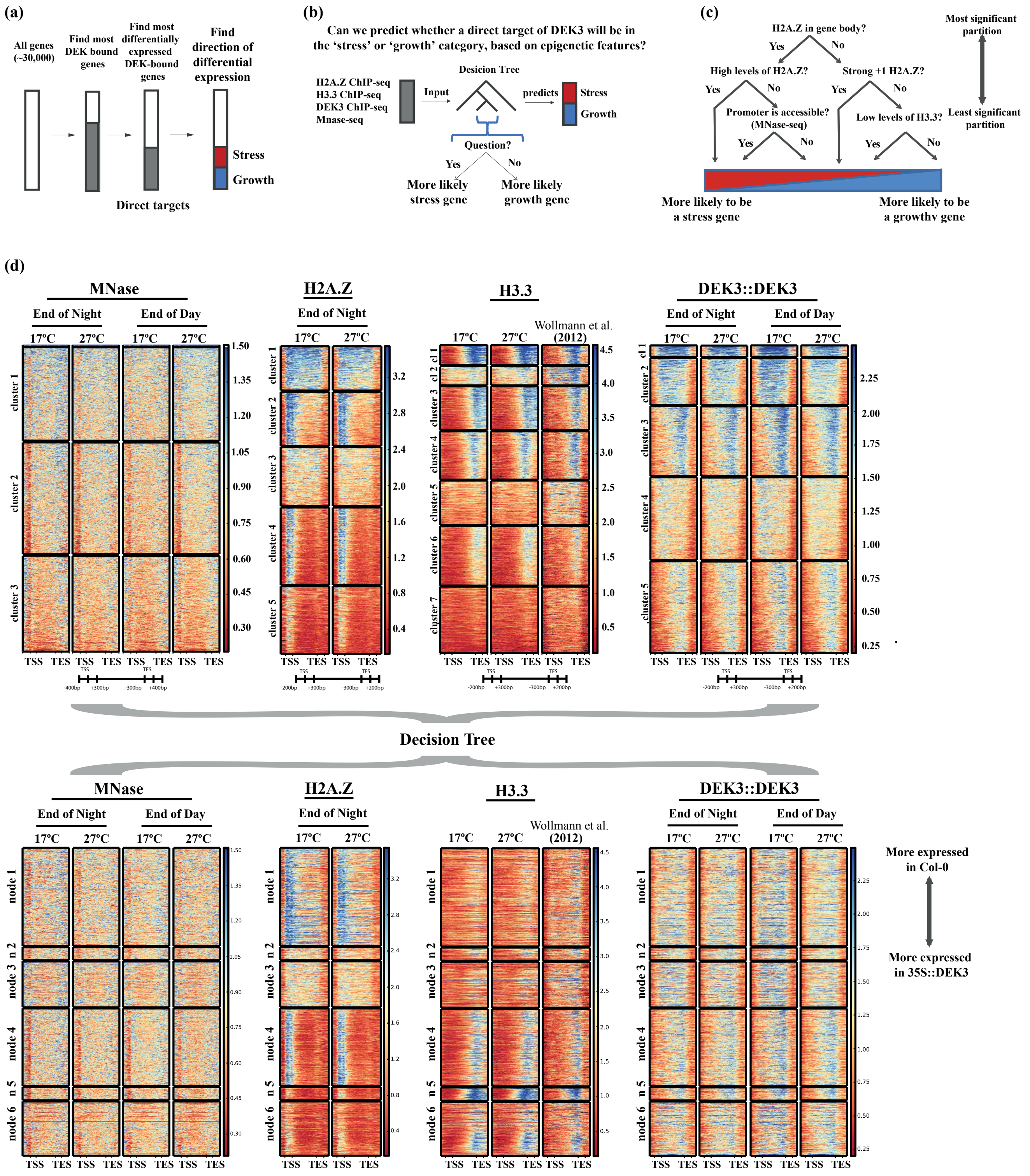

### Figure 3 supp

(e)

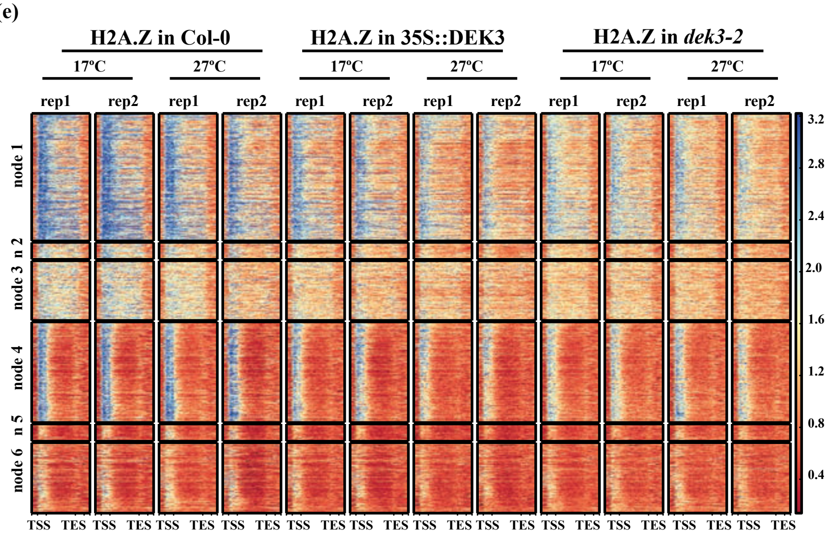


(f)

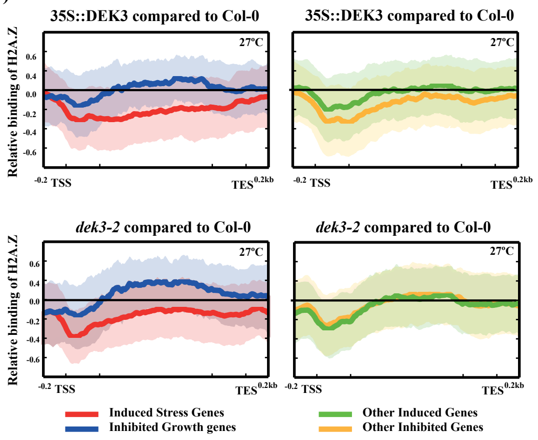

#### Supplementary Fig. 4.

#### DEK3 controls the expression of “growth” and “stress” genes in similar way to stress activation and the antagonistic relationship between DEK3 and H2A.Z may be conserved

(a) This depicts the log-fold change in the DEK3 target gene expression acquired from the AtGeneExpress (compared to a control non-stress treated samples). The genes are organised in the same order as in Figure 2c.
(b) Expression of “Growth” and “Stress” genes in Col-0, *dek3-2* and *35S::DEK3* genetic backgrounds based on the z-scores calculated from the transcript per million (TPM). Plants has been grown at 17°C and 27°C, and collected at 8 time points during 24 h. The lines represent the best fitting line for each set of samples with 95% confidence intervals in gray. Black bars on the top indicate night, white bars day.
(c) Acceleration of flowering of Col-0, *dek3-2*, *arp6-1* and *dek3-2 arp6-1* plants grown under short day photoperiod at 17°C, 22°C and 27°C.
(d) Boxplots summarising leaf number at the time of flowering of the plants from (c). The numbers of plants used: (17/22/27°C) Col-0 12/12/12; *dek3-2* 12/12/12; *arp6-1* 10/12/12; *dek3-2 arp6-1* 12/10/12. Box and whisker plots show median, inter-quartile ranges and 95% confidence intervals (t-test p-values are summarized in Supp. Table 5).
(e) Boxplots summarising days to bolting of the plants from (c). The numbers of plants used: (17/22/27°C) Col-0 12/12/12; *dek3-2* 12/12/12; *arp6-1* 12/12/12; *dek3-2 arp6-1* 12/12/12. Box and whisker plots show median, inter-quartile ranges and 95% confidence intervals (t-test p-values are summarized in Supp. Table 5).
(f) Boxplots summarising the hypocotyl length of Col-0, *dek3-2*, *35S::DEK3_1* and *35S::DEK3_2* (independent over-expression lines. Seedlings have been grown for 7 days under short day photoperiod at 17°C, 22°C and 27°C. The numbers of seedlings used: (17/22/27°C) Col-0 35/39/32; *dek3-2* 33/36/35; *arp6-1* 27/36/37; *dek3-2 arp6-1* 44/41/36. Box and whisker plots show median, inter-quartile ranges and 95% confidence intervals (t-test p-values are summarized in Supp. Table 5).
(g) Expression of “Growth” and “Stress” genes in Col-0, *arp6-1*, *dek3-2* and *dek3-2 arp6-1* genetic backgrounds were hierarchically clustered, based on the log2 ratio compared to the values of Col-0 in transcript per million (TPM). Plants has been grown at 17°C and 27°C and collected at the end of the day (white bar) or at the end of the night (black bar). Up-regulated genes are shown in red and down-regulated genes are shown in blue. The sidebar on the left of the heatmap indicates the major clusters.
(h) *35S::DEK3 arp6* plants show temperature dependent lethality when grown in different temperatures (17°C, 22°C and 27°C).
(i) The model of DEK3 regulation of the trade-off between “growth” and “arrest” genes via H2A.Z-nucleosomes in Arabidopsis in different environmental conditions. The “growth” genes (upper

panel) and “stress” genes (lower panel) are distinguished mainly by their gene-body distribution of H2A.Z. While “stress” genes have high gene body H2A.Z occupancy, which will be lost by over-expression of DEK3 allowing their induction in response to environmental stimuli only in this genotype; “growth” genes have predominantly +1 H2A.Z nucleosomes and will acquire gene-body H2A.Z in the absence of DEK3, causing their induction. Chromatin landscape is likely to be changing as a result of DEK3-H2A.Z interaction.

(j) Summary of the known hDEK up- and down-regulated genes based on literature.

(k) Representative profiles based on published genome-wise H2A.Z binding[2] to the published DEK target genes in humans [3–11].


1. Simon, W. Differential expression analysis for sequence count data. *Genome Biol.* **11,** (2010).
2. Sequeira-Mendes, J. *et al.* The Functional Topography of the Arabidopsis Genome Is Organized in a Reduced Number of Linear Motifs of Chromatin States. *Plant Cell* **26,** 2351–2366 (2014).
3. Koleva, R. I. *et al.* HEMATOPOIESIS AND STEM CELLS C / EBP ␣ and DEK coordinately regulate myeloid differentiation. **119,** 1–3 (2012).
4. Sammons, M. *et al.* Negative regulation of the RelA/p65 transactivation function by the product of the DEK proto-oncogene. *J. Biol. Chem.* **281,** 26802–26812 (2006).
5. Kim, K.-B. *et al.* Negative regulation of peroxiredoxin 6 (Prdx 6) transcription by nuclear oncoprotein DEK during leukemia cell differentiation. *Animal Cells Syst. (Seoul).* **18,** 318–323 (2014).
6. Hu, H. G., Illges, H., Gruss, C. & Knippers, R. Distribution of the chromatin protein DEK distinguishes active and inactive CD21/CR2 gene in pre- and mature B lymphocytes. *Int. Immunol.* **17,** 789–796 (2005).
7. Zhang, Y. *et al.* The DEK oncogene activates VEGF expression and promotes tumor angiogenesis and growth in HIF-1 a -dependent and -independent manners. **7,** (2016).
8. Campillos, M., García, M. A., Valdivieso, F. & Vázquez, J. Transcriptional activation by AP-2α is modulated by the oncogene DEK. *Nucleic Acids Res.* **31,** 1571–1575 (2003).
9. Karam, M. *et al.* Chromatin Redistribution of the DEK Oncoprotein Represses hTERT Transcription in Leukemias. *Neoplasia* **16,** 21-W14 (2014).
10. Hu, H. gang, Scholten, I., Gruss, C. & Knippers, R. The distribution of the DEK protein in mammalian chromatin. *Biochem. Biophys. Res. Commun.* **358,** 1008–1014 (2007).
11. Lohmann, F. *et al.* The DEK Oncoprotein Is a Critical Component of the EKLF / KLF1 Enhancer in Erythroid Cells. **35,** 3726–3738 (2015).

### Figure 4 supp

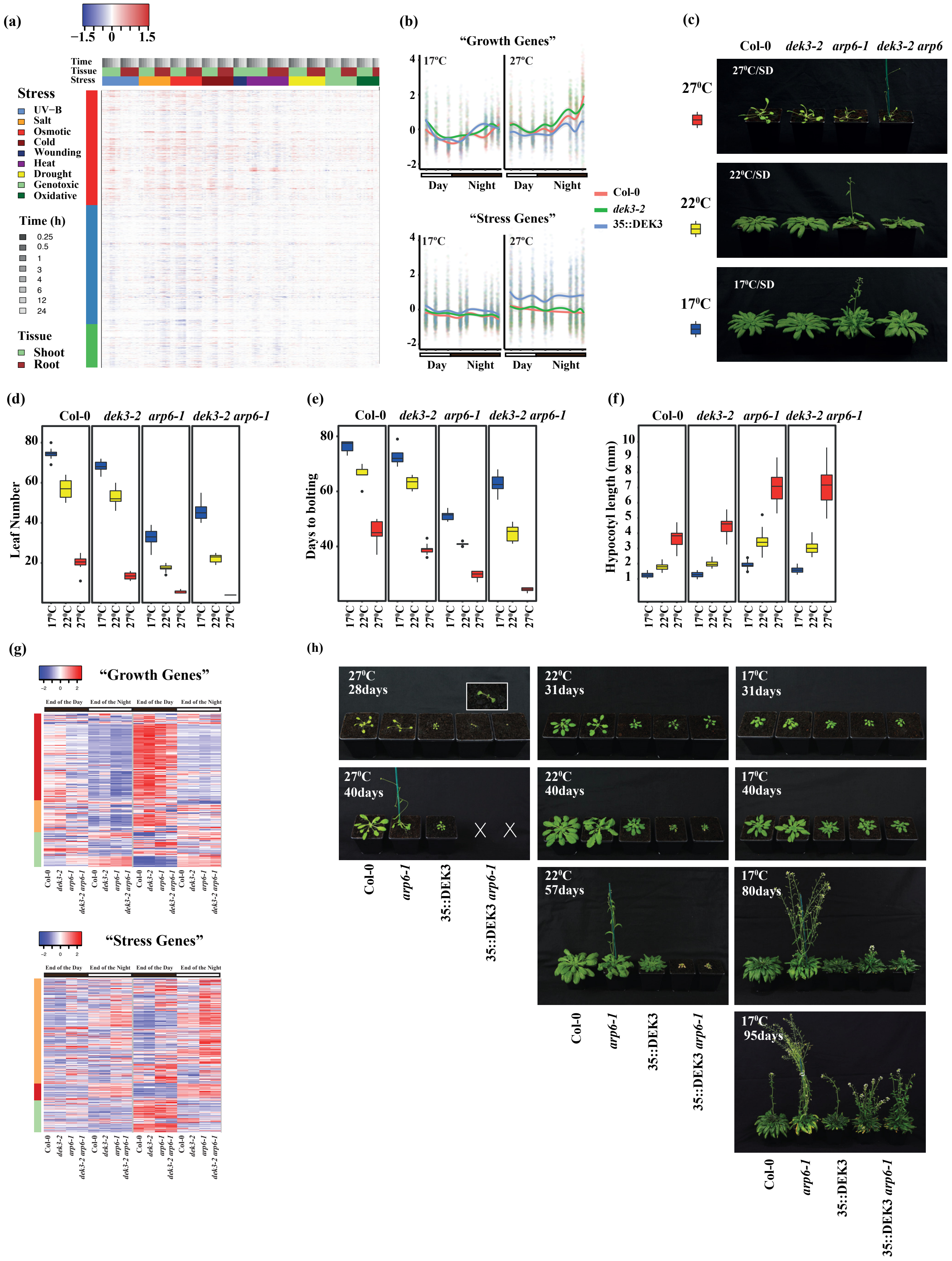

### Figure 4 supp_page2

(i)

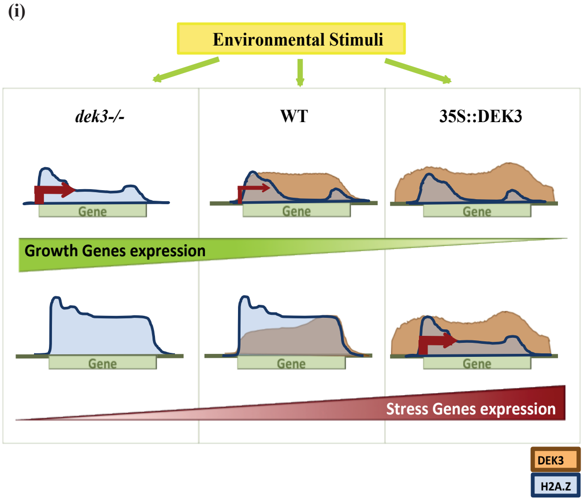


(j)

| Up-regulated genes by hDEK | Down-regulated genes by hDEK |
|---|---|
| KLF1 | PRDX6 |
| APOE | RELA |
| CEBPE | CR2 |
| VEGFA | TERT |
| CSF3R | |
| TOP1 | |

(k)

**H2A.Z binding profiles in MCF7 cells**

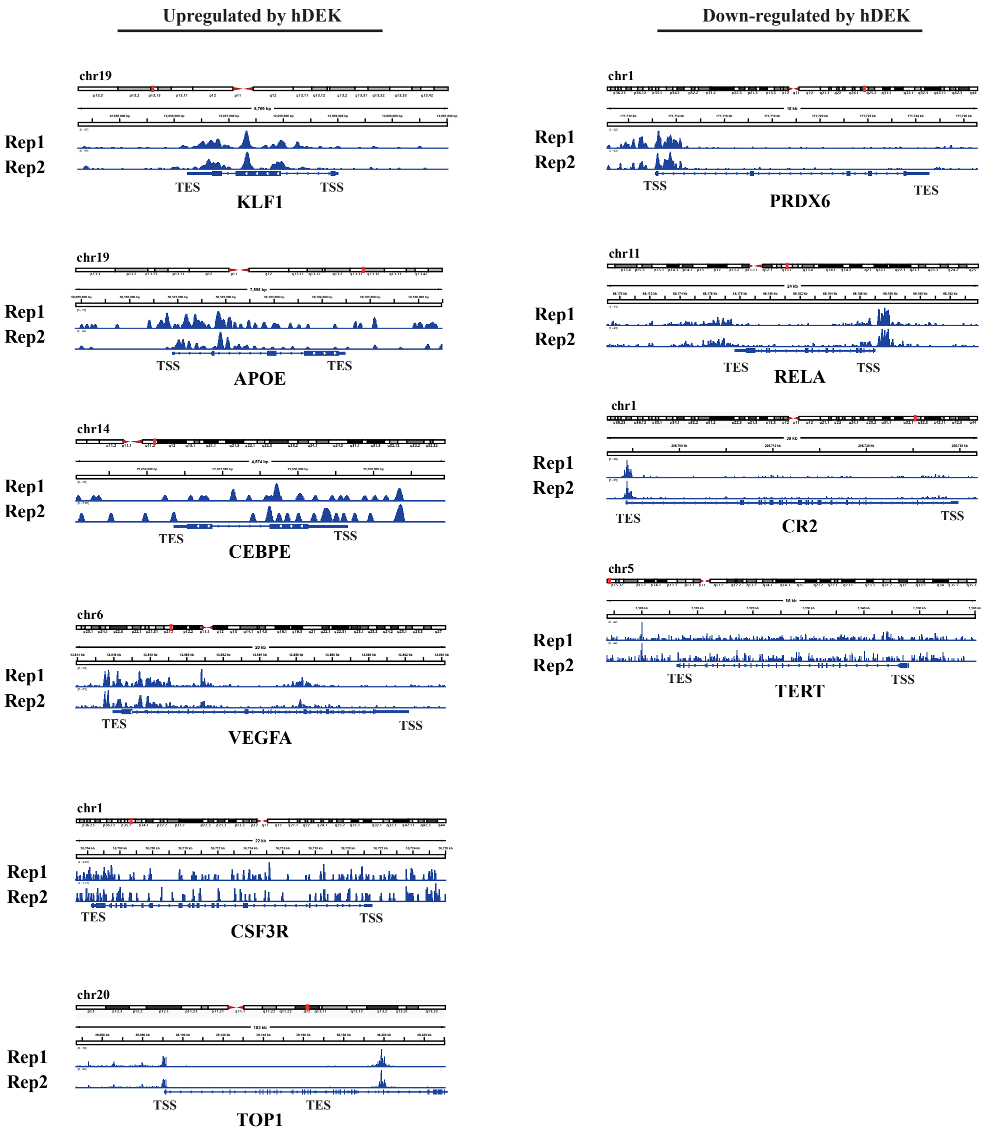
